# The regulatory protein ChuP connects heme and siderophore-mediated iron acquisition systems required for *Chromobacterium violaceum* virulence

**DOI:** 10.1101/2022.02.10.479982

**Authors:** Vinicius M. de Lima, Bianca B. Batista, José F. da Silva Neto

## Abstract

*Chromobacterium violaceum* is an environmental Gram-negative beta-proteobacterium that causes systemic infections in humans. *C. violaceum* uses siderophore-based iron acquisition systems to overcome the host-imposed iron limitation, but its capacity to use other iron sources is unknown. In this work, we characterized ChuPRSTUV as a heme utilization system employed by *C. violaceum* to explore an important iron reservoir in mammalian hosts, free heme and hemoproteins. We demonstrate that the *chuPRSTUV* genes comprise a Fur-repressed operon that is expressed under iron limitation. The *chu* operon potentially encodes a small regulatory protein (ChuP), an outer membrane TonB-dependent receptor (ChuR), a heme degradation enzyme (ChuS), and an inner membrane ABC transporter (ChuTUV). Our nutrition growth experiments using *C. violaceum chu* deletion mutants revealed that, with the exception of *chuS*, all genes of the *chu* operon are required for heme and hemoglobin utilization in *C. violaceum*. The mutant strains without *chuP* displayed increased siderophore halos on CAS plate assays. Significantly, we demonstrate that ChuP connects heme and siderophore utilization by acting as a positive regulator of *chuR* and *vbuA*, which encode the TonB-dependent receptors for the uptake of heme (ChuR) and the siderophore viobactin (VbuA). Our data favor a model of ChuP as a heme-binding post-transcriptional regulator. Moreover, our virulence data in a mice model of acute infection demonstrate that *C. violaceum* uses both heme and siderophore for iron acquisition during infection, with a preference for siderophores over the Chu heme utilization system.

## INTRODUCTION

Iron is an essential micronutrient required as a cofactor of proteins involved in different cellular processes (Braun and Hantke, 2011; Palmer and Skaar, 2016). The ability to vary from soluble ferrous (Fe^2+^) to insoluble ferric (Fe^3+^) states confers iron its catalytic properties but can result in high toxicity and low bioavailability (Braun and Hantke, 2011; Huang and Wilks, 2017). As a metal essential for both hosts and pathogens, iron is at the center of an evolutionary battle (Skaar, 2010; Hood and Skaar, 2012; Parrow et al., 2013; Sheldon et al., 2016). Hosts restrict iron availability using iron-sequestering proteins like transferrin, lactoferrin, haptoglobin, hemopexin, and calprotectin, a process known as nutritional immunity (Hood and Skaar, 2012; Cassat and Skaar, 2013; Ganz and Nemeth, 2015). Conversely, pathogens subvert the host-imposed iron limitation by employing strategies such as the production, release, and uptake of low-molecular-weight iron chelators (siderophores such as enterobactin) and high-affinity heme-binding proteins (hemophores such as HasA) (Wandersman and Delapelaire, 2012; Runyen-Janecky et al., 2013; Contreras et al., 2014; Sheldon et al., 2016).

Heme is a tetrapyrrole that coordinates iron (Fe^2+^) at its center. It is a cofactor of proteins like cytochromes and catalases. Therefore, almost every organism requires heme, which is obtained by synthesis and/or uptake from exogenous sources (Runyen-Janecky, 2013; Choby and Skaar, 2016). The greatest iron reservoir in mammals is the heme bound into hemoglobin found inside the erythrocytes. Many bacteria use heme and hemoproteins as an iron source, and the preference for heme or siderophore as the main iron acquisition strategy varies according to the bacterium and the infection status (Runyen-Janecky, 2013; Choby and Skaar, 2016; Sheldon et al., 2016; Zygiel et al., 2020). Heme uptake/utilization systems have been described in several bacterial pathogens, including *Pseudomonas aeruginosa* (Has, Phu, and Hxu), *Yersinia* spp (Hem and Hmu), *Escherichia coli* (Chu), and *Staphylococcus aureus* (Isd). In Gram-negative bacteria, the import of heme involves high-affinity TonB-dependent receptors (TBDRs) in the outer membrane (e.g., PhuR) and ABC-type transport systems in the periplasm and inner membrane (e.g., PhuTUV) (Eakanunkul et al., 2005; Noinaj et al., 2010; Fournier et al., 2011; Choby and Skaar, 2016; Huang and Wilks, 2017; Klebba et al., 2021). Once in the cytosol, heme is degraded by canonical or non-canonical heme oxygenases, releasing iron and other compounds (Contreras et al., 2014; Lamattina et al., 2016).

Genes encoding heme uptake systems are under complex regulation. They are regulated by Fur, a metalloregulator that uses Fe^2+^ as cofactor to repress the expression of iron uptake systems (da Silva Neto et al., 2009; Chandrangsu et al., 2017; Sarvan et al., 2018) and activated by heme-dependent regulatory systems, such as the extracytoplasmic function (ECF) sigma factor signaling cascade Has (Wandersman and Delapelaire, 2012; Huang and Wilks, 2017). Small proteins from the HemP/HmuP family have been described as required for heme utilization by regulating the expression of heme uptake genes. However, the proposed regulatory mechanisms are quite distinct. In *Bradyrhizobium japonicum* and *Burkholderia multivorans*, the HemP/HmuP proteins were described as direct transcriptional activators (Escamila-Hernandez and O’Brian, 2012; Sato et al., 2017), while in *Ensifer meliloti* (formerly *Sinorhizobium meliloti*), HmuP appears to act as a post-transcriptional activator (Amarelle et al., 2010; Amarelle at al., 2019).

*Chromobacterium violaceum* is a Gram-negative beta-proteobacterium found in the water and soil of tropical and subtropical regions that causes opportunistic human infections with high mortality rates (Yang and Li, 2011; Kumar, 2012; Khalifa et al., 2015; Batista and da Silva Neto, 2017). An important virulence determinant in *C. violaceum* is the Cpi1/1a type III secretion system involved in hepatocyte invasion and innate immune system activation (Miki et al., 2010; Zhao et al., 2011; Maltez et al., 2015). Recently, we demonstrated that *C. violaceum* relies on the regulator Fur, two putative endogenous catecholate-type siderophores, and the siderophore-acquisition TBDRs CbuA and VbuA to overcome host-imposed iron limitation (Batista et al., 2019; Santos et al., 2020). However, *C. violaceum* mutants lacking siderophores had moderate attenuation in virulence in a mouse model of acute infection (Batista et al., 2019), suggesting that *C. violaceum* uses siderophore-independent mechanisms for iron acquisition during infection. In the current work, we demonstrate that an operon with six genes, here named *chuPRSTUV* (*chu* – *<u>C</u>hromobacterium* <u>h</u>eme <u>u</u>tilization), encodes a Fur-regulated heme uptake system (ChuRTUV) that is required for heme and hemoglobin utilization in *C. violaceum*. We also show that the small heme-binding protein ChuP is required for heme and siderophore-mediated iron acquisition by acting as a post-transcriptional activator of the TBDR genes *chuR* and *vbuA*. Furthermore, using *in vivo* virulence assays in mice, we demonstrate that these heme and siderophore-mediated iron uptake systems work together to help *C. violaceum* overcome iron limitation in the host.

## MATERIALS AND METHODS

### Bacterial strains, plasmids, and growth conditions

The bacterial strains and plasmids used in this work are indicated in Table 1. *E. coli* strains were cultured in Luria-Bertani (LB) medium at 37 °C. *C. violaceum* strains were cultured in LB medium or M9 minimal medium supplemented with 0.1% casein hydrolysate (M9CH) at 37 °C (Batista et al., 2019). The cultures were supplemented with kanamycin (50 μg/mL), tetracycline (10 µg/mL), or ampicillin (100 μg/mL), when necessary. Iron deficiency was obtained by the addition of 2,2’-dipyridyl (DP) (Sigma) to the medium, while iron sufficiency was achieved by supplementation with FeSO_4_ (Sigma), hemin (Hm) (Sigma), or hemoglobin (Hb) (Sigma).

**Table 1.**
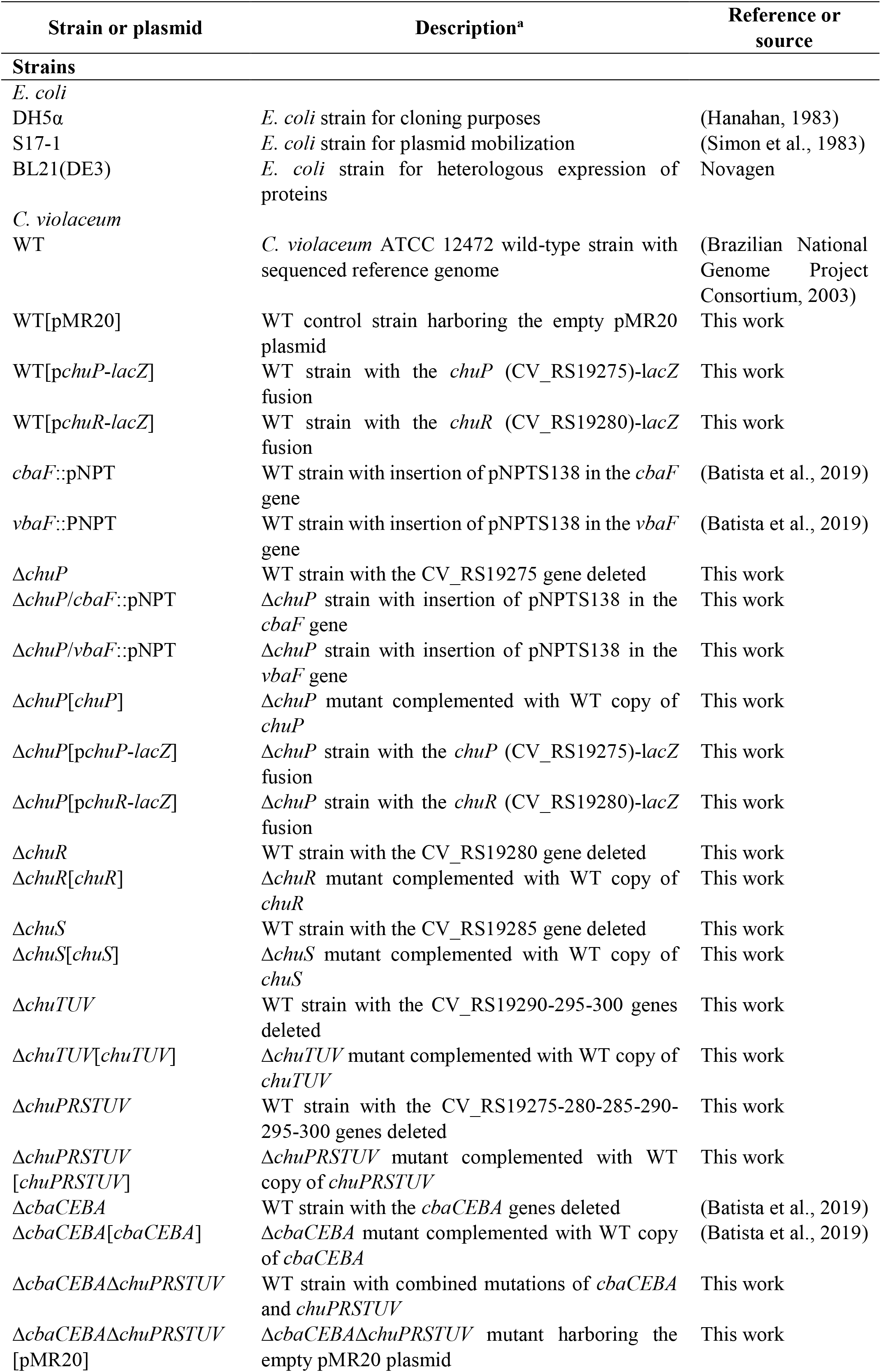
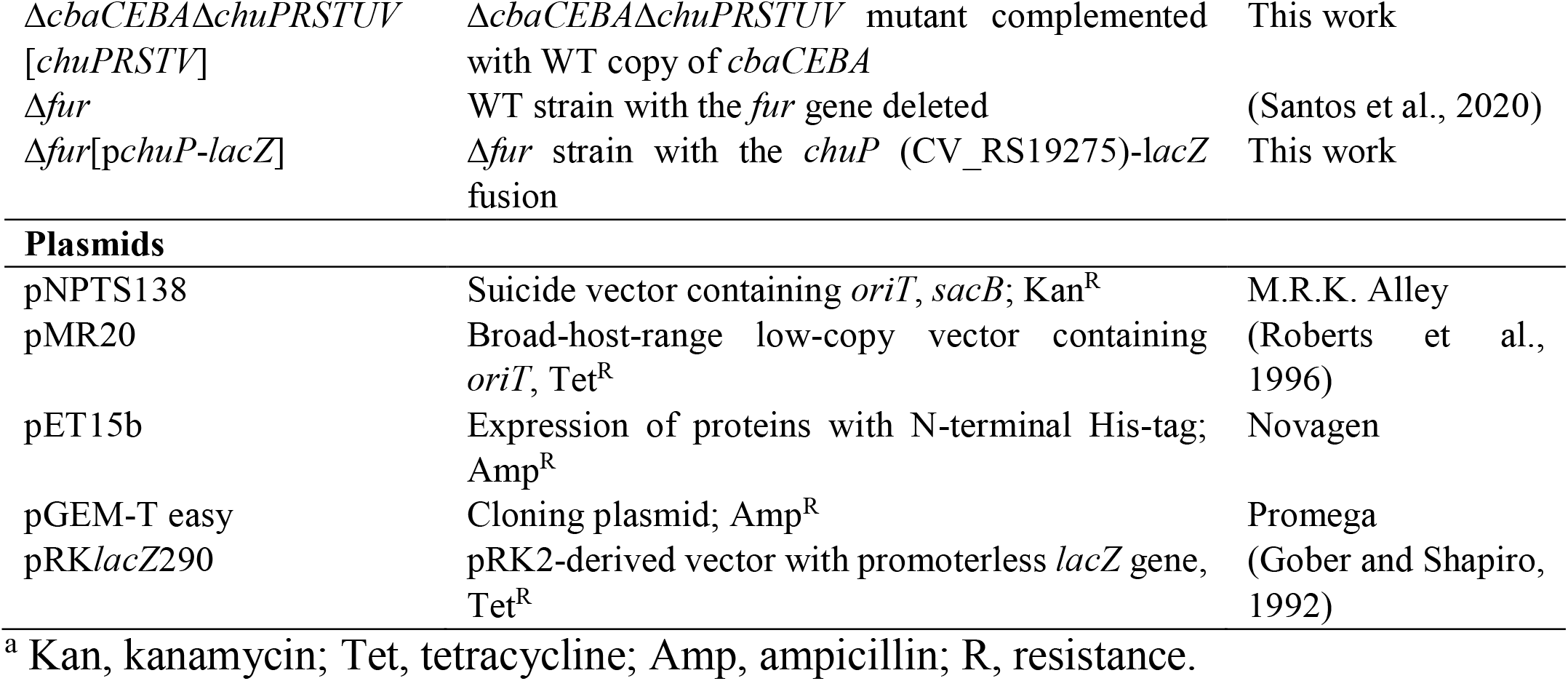
Bacterial strains and plasmids.

### Construction of *C. violaceum* mutant and complemented strains

Null mutant strains were generated by a previously established allelic exchange mutagenesis protocol (da Silva Neto et al., 2012; Batista et al., 2019; Santos et al., 2020). In-frame null-deletion mutants derived from the wild-type *C. violaceum* ATCC 12472 strain, with the exception of the Δ*cbaCEBA*Δ*chuPRSTUV* strain that was obtained using the Δ*cbaCEBA* mutant as background (Batista et al., 2019). The insertion mutants for the non-ribosomal peptide synthetase (NRPS) genes *cbaF* and *vbaF* were obtained by a protocol based on a single recombination event (Batista et al., 2019). For genetic complementation, the *chuP, chuR, chuS, chuTUV*, and *chuPRSTUV* genes were amplified by PCR, cloned into the low-copy-number plasmid pMR20, and transferred to the mutant strains by conjugation. The primers used for cloning, sequencing, and mutant confirmation are listed in Table S1.

### MIC assay

To achieve iron-limited conditions in M9CH for *C. violaceum*, we determined the minimal inhibitory concentration (MIC) of DP in this medium, as previously performed in LB medium (Batista et al., 2019). Wild-type *C. violaceum* overnight cultures were diluted to an optical density at 600 nm (OD_600_) of 0.01 in M9CH, without or with DP (100 µM, 112.5 µM, 125 µM, 132.5 µM, and 150 µM), and grown under agitation (250 rpm) at 37 °C. The MIC of 132.5 µM DP for the WT strain was established based on the turbidity of the cultures after 24 h cultivation.

### Growth curves

Growth curves were determined in M9CH without or with Hm. Overnight cultures of *C. violaceum* strains were diluted in 5 mL of M9CH to an OD_600_ of 0.02. Then, a new dilution (1:2) was performed to achieve an OD_600_ 0.01 and the required concentrations of Hm in 200 µL final M9CH in 96-well plates. The plates were incubated at 37 °C under moderate orbital agitation in SpectraMax i3 MiniMax Imaging Cytometer (Molecular Devices). The measurements of OD_600_ were recorded every 15 minutes over 24 hours. The experiment was performed in three biological replicates.

### Heme and hemoglobin nutrition assay

The ability of *C. violaceum* to use Hm and Hb as iron sources was assessed using a nutrition assay (Balhesteros et al., 2017) with some modifications. *C. violaceum* overnight cultures in M9CH were diluted to an OD_600_ of 1.0 in M9CH. Then, 25 µL of each dilution were embedded in 25 mL of iron-depleted M9CH 0.8% agar (containing 50 µM, 100 µM, 125 µM or 150 µM of DP). Paper discs were added onto the plate surface, and 10 µL aliquots of 100 µM Hm, 20 mM NaOH, 150 µM Hb, and 100 mM NaCl were applied to individual discs. After incubation for 16 h at 37 °C, we inspected for growth halos that developed around the discs. The growth area was quantified using the Image J software and normalized by subtracting the disc areas. The experiment was performed in three biological replicates.

### Cell viability in the presence of heme

The toxic concentrations of Hm for *C. violaceum* were assessed by cell viability. Overnight cultures were diluted to an OD_600_ of 0.01 in M9CH without or with Hm (30 µM, 600 µM, and 2000 µM), and grown under agitation (250 rpm) for 24 h at 37 °C. Serial dilution in phosphate-buffered saline (PBS) was performed, and 10 µL were spotted onto M9CH plates. Hemin toxicity was determined based on the colony-forming units (CFU) displayed by the strains after incubation for 24 h at 37 °C. These experiments were performed in three biological replicates.

### Hemolysis assay

The hemolytic activity was assessed in 5% (v/v) sheep-blood Mueller-Hinton agar plates. Five microliters of *C. violaceum* M9CH overnight cultures were spotted onto the plate. The hemolytic activity was detected by the lighter halos that developed due to erythrocyte lysis after incubation for 7 days at 37 °C. The area of activity was quantified using the Image J software and normalized by subtracting the bacterial growth area. The experiment was performed in three biological replicates.

### Siderophore activity assay

Siderophore activity was detected by chrome azurol S (CAS) plate assay in modified peptone-sucrose agar (PSA-CAS) plates (Batista et al., 2019; Santos et al., 2020). Ten microliters of *C. violaceum* overnight cultures in M9CH were spotted onto the plate surface, and the siderophore activity was determined by the orange halos that developed after incubation for 24 hours at 37 °C. The area of activity was quantified using the Image J software and normalized by subtracting the bacterial growth area. The experiment was performed in three biological replicates.

### Transcriptional *lacZ* fusions and β-galactosidase assays

The upstream regions of genes of interest were amplified by PCR with specific primers (Table S1) and cloned into the pGEM-T easy plasmid (Promega). After digestion with proper restriction enzymes (Table S1), the inserts were subcloned into the pRKl*acZ*290 vector to generate transcriptional fusions to the *lacZ* gene. *C. violaceum* cultures harboring the reporter plasmids were grown until an OD_600_ of 0.6 – 0.8 in M9CH, and either untreated or treated with 100 µM Hm or 100 µM FeSO_4_ for 2 h. Bacterial cells were assayed for β-galactosidase activity as previously described (Santos et al., 2020). The experiment was performed in three biological replicates.

### Co-transcription by RT-PCR

The *C. violaceum* wild-type strain was grown in M9CH until an OD_600_ of 1.0 – 1.2. Total RNA was extracted using Trizol reagent (Invitrogen) and purified with Direct-zol™ RNA Miniprep Plus (Zymo Research). RT-PCR was performed with the SuperScript III One-Step RT-PCR System with Platinum Taq High Fidelity DNA Polymerase (Invitrogen). One microgram of each RNA sample and specific primers (Table S1) that amplify regions from *chuP* to *chuR* (439 bp), *chuR* to *chuS* (373 bp), and *chuS* to *chuT* (662 bp) were used in the reactions. PCRs using conventional Taq DNA polymerase, and the same sets of primers, were performed with genomic DNA (positive control) and RNA (negative control) as templates.

### Gene expression by RT-qPCR

The *C. violaceum* wild type, Δ*chuP*, and Δ*chuP*[*chuP*] strains were grown in M9CH until midlog growth phase, and the cultures were either untreated or treated with 100 µM Hm or 100 µM FeSO_4_ for 2 h. Total RNA was extracted and purified as described above. Two micrograms of total RNA from each sample were converted to cDNA using the High-Capacity cDNA Reverse Transcription kit (Thermo Fisher Scientific). Genomic DNA contamination (for RNA) and reverse transcription efficiency (for cDNA) were checked by conventional PCR with the primers for the *rpoH* gene (Table S1). Quantitative PCR (qPCR) reactions were performed using the PowerUp™ SYBR™ Green Master Mix (Thermo Fisher Scientific), the specific primers (Table S1), and 0.5 µL of cDNA. The relative expression was calculated by the 2^-ΔΔ^_Ct_ method (Livak and Schmittgen, 2001). Data from three biological replicates were normalized by an endogenous control (*rpoH* gene) and a reference condition (WT in M9CH 100 µM Hm).

### Expression and purification of the recombinant ChuP

The coding region of *chuP* was amplified by PCR (Table S1) and cloned into the pET-15b plasmid (Table 1). After induction in *E. coli* BL21(DE3) with 1 mM Isopropyl *β*-D-1-thiogalactopyranoside (IPTG) for 2 h at 37 °C, the His-ChuP protein was purified from the soluble extract by affinity chromatography in a Ni-NTA Superflow column (Qiagen). The elution fractions were evaluated using 18% SDS-PAGE. The aliquots containing the purified His-ChuP were concentrated using a VivaSpin 6 column (Sartorius), and desalted by gel filtration in PD-10 column (GE Healthcare) in storage buffer (100 mM NaH_2_PO_4_, 600 mM NaCl, 20% glycerol, pH 8) (Puri and O’Brien, 2006). The concentration of the His-ChuP protein was determined by measurement of OD at 280 nm and using its extinction coefficient calculated by the Protparam Tool (ExPASy) (http://web.expasy.org/protparam).

### Heme binding assay

The ability of the recombinant His-ChuP protein to interact with Hm was evaluated by spectrophotometry (Puri and O’Brien, 2006; Amarelle et al., 2016). The binding reactions were performed in interaction buffer (50 mM Na_2_HPO_4_, 300 mM NaCl, 10% glycerol, pH 8), using 10 µM of His-ChuP and a range of Hm concentrations for 5 minutes at 25 °C in the dark. After incubation, the absorbance between the wavelengths 300 and 600 nm was measured with 10 nm increments on a SpectraMax i3 MiniMax Imaging Cytometer. The binding of ChuP to Hm was determined by the change in absorbance at 413 nm fit to one-site binding model non-linear regression on Graph Pad Prism 7.

### Electrophoretic mobility shift assay (EMSA)

DNA sequences upstream of *chuP, chuR*, and CV_2599 were amplified by PCR using the primers listed in Table S1. The DNA fragments were radiolabeled and used for interaction with His-ChuP following a previously described protocol (da Silva Neto et al., 2009; Previato-Mello et al., 2017), with the modification of adding the CV_2599 promoter fragment (negative control) in the same reaction.

### Mouse virulence assays

Virulence assays were performed in a mouse intraperitoneal (i.p.) model of *C. violaceum* infection as previously established (Previato-Mello et al., 2017; Batista et al., 2019). Bacterial strains were diluted to an OD_600_ of 0.01 and cultured in 5 mL LB for 20 h at 37 °C. A dose of 10^6^ CFU in PBS was injected into 6-week-old female BALB/c mice, and the animals were monitored for 7 days post-infection. To assess the bacterial burden in the liver and spleen, mice were infected as above and euthanized 20 h or 96 h post-infection (h.p.i.). The organs were aseptically collected, homogenized in PBS, and the dilutions were plated for CFU counting. Mice were obtained and maintained at the Animal Facilities of Ribeirão Preto Medical School (FMRP-USP). The assays were performed according to the Ethical Principles in Animal Research adopted by the National Council for the Control of Animal Experimentation (CONCEA). The animal ethics protocol 146/2019 was approved by the Local Ethics Animal Committee (CEUA) of FMRP-USP.

### Statistical analysis

Data collected were employed for statistical analysis in GraphPad Prism version 7. For the column graphs, the normality test was performed using Shapiro-Wilk’s test. Statistically significant p values and the tests that were performed are indicated in the figure legends.

## RESULTS

### The *chuPRSTUV* operon is regulated by Fur according to the iron levels

*In silico* analysis of the *C. violaceum* ATCC 12472 genome sequence revealed a gene cluster with six genes (CV_RS19275-280-285-290-295-300) that resembles an operon encoding a putative heme utilization system. These genes, here named *chuPRSTUV*, are annotated as a HemP/HmuP family regulator (ChuP), a TonB-dependent receptor (ChuR), a hemin degrading factor (ChuS), and an ABC-transport system (ChuTUV) (Figure 1A). To evaluate if the *chuPRSTUV* genes are organized into an operon, we performed RT-PCR reactions using RNA from the WT strain grown in M9CH and a set of primers that amplify regions between *chuPR, chuRS*, and *chuST* genes (Figure 1A, B). After reverse transcription and amplification, bands with the expected sizes were detected for the three tested primer combinations, confirming that the *chuPRSTUV* genes are indeed co-transcribed (Figure 1B).

**Figure 1.**
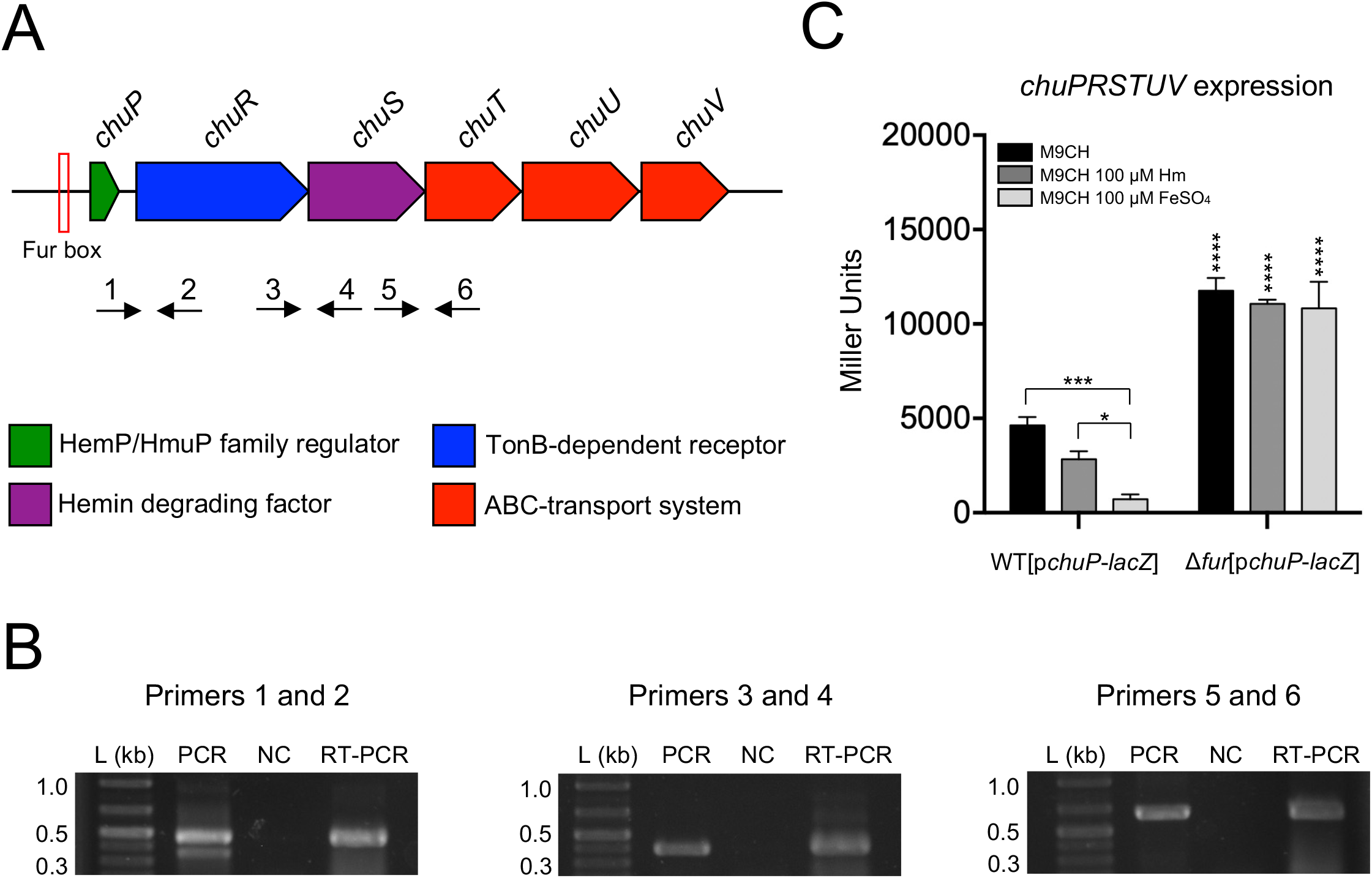
The *chuPRSTUV* genes compose an operon regulated by the iron levels and Fur. **(A)** Genomic organization of the *chuPRSTUV* genes in *C. violaceum*. A predicted Fur box is indicated. Numbered arrows indicate the primers used in RT-PCR (not scaled). **(B)** Confirmation of co-transcription of the *chuPRSTUV* genes. The RT-PCR reactions amplified fragments of 439 bp (Primers 1 and 2), 373 bp (Primers 3 and 4), and 662 bp (Primers 5 and 6). Conventional PCR was performed using genomic DNA (PCR) and RNA (NC) as controls. L, 1 Kb plus DNA Ladder (Thermo Scientific). **(C)** Promoter activity of the *chu* operon in response to iron and Fur. *β*-galactosidase assays were performed from the WT and Δ*fur* strains harboring the *chuP-lacZ* fusion grown in M9CH medium and either untreated or treated with 100 *μ*M Hm or 100 *μ*M FeSO_4_. Data are from three biological replicates. ****, p < 0.0001; ***, p < 0.001; *, p < 0.05; when not shown, n.s. (not significant). Two-way ANOVA followed by Dunnett’s multiple-comparison test.

Our inspection of the promoter region of *chuP* revealed a putative Fur binding site sequence (ATGATAATGGTTATCATT) that resembles Fur boxes found in other bacteria (Sarvan et al., 2018). To investigate whether the *chu* operon is regulated by iron and Fur, we cloned the promoter region of *chuP* into a *lacZ* reporter plasmid. The WT and Δ*fur* strains harboring the p*chuP*-*lacZ* fusion were used to assess the *chuP* promoter activity by *β*-galactosidase assay in M9CH medium, which was previously reported as an iron-limited condition (Santos et al., 2020) and under iron sufficiency (M9CH supplemented with Hm or FeSO_4_) (Figure 1C). The promoter activity was higher under iron-limited (M9CH) than iron-replete conditions in the WT strain. Interestingly, the reduction in activity was higher with FeSO4 than with Hm supplementation. In the Δ*fur* mutant, the promoter was highly active regardless of iron levels. Moreover, the activity was twofold higher than that detected for the WT strain in M9CH, suggesting total promoter de-repression in the absence of Fur (Figure 1C). Altogether, these results demonstrate that the *chuPRSTUV* genes comprise a Fur-repressed operon that is expressed under iron limitation.

### The *chuPRSTUV* operon encodes a heme uptake system (ChuRTUV) and a regulatory protein (ChuP) required for heme and hemoglobin utilization

To characterize the role of the *chuPRSTUV* operon in *C. violaceum*, we generated null-mutant strains deleted for single genes (Δ*chuP*, Δ*chuR*, and Δ*chuS*) or multiple genes (Δ*chuTUV* and Δ*chuPRSTUV*) of the *chu* operon, and their respective complemented strains. We also obtained a mutant strain lacking both the *chu* operon and the *cbaCEBA* genes (Δ*cbaCEBA*Δ*chuPRSTUV*). The CbaCEBA enzymes are involved in the synthesis of 2’3-DHB, the precursor of catecholate-type siderophores in *C. violaceum* (Batista et al., 2019). All mutants showed regular fitness, as assessed by growth curves in M9CH and M9CH plus heme and by cell viability in LB (Supplementary Figure 1).

To test the involvement of the *C. violaceum chu* genes in heme and hemoglobin utilization, we developed a nutrition assay providing 100 µM Hm or 150 µM Hb as alternative iron sources in DP-iron chelated M9CH medium (Figure 2). To visualize the growth halos, we adjusted the DP concentrations as the lowest able to avoid confluent growth for each strain (50 µM for Δ*cbaCEBA*, 100 µM for Δ*cbaCEBA*Δ*chuPRSTUV*, 125 µM for WT, Δ*chuP*, Δ*chuR*, Δ*chuS*, and Δ*chuPRSTUV*, and 150 µM for Δ*chuTUV*). Under these conditions, the WT and the Δ*chuS* strains formed Hm and Hb-stimulated growth halos. All the other mutant strains of the *chuPRSTUV* operon lost the ability to grow when Hm and Hb were provided as iron sources (Figure 2). For the Δ*chuR* strain, a very weak growth stimulus could still be detected only in the presence of heme (Figure 2B). As expected, the Δ*cbaCEBA* mutant that does not synthesize siderophores grew similarly to the WT strain after supplementation with Hm and Hb. Genetic complementation of the mutant strains fully restored the growth in Hm and Hb under deficiency with 125 µM DP (Figure 2). Taken together, these results demonstrate that the *chuPRSTUV* operon encodes a heme uptake system (ChuRTUV) that is also involved in hemoglobin utilization. Moreover, the weak growth detected for the Δ*chuR* mutant with Hm but not with Hb suggests that *C. violaceum* has additional mechanisms in the outer membrane for heme uptake but relies specifically on ChuR for heme uptake from hemoglobin.

**Figure 2.**
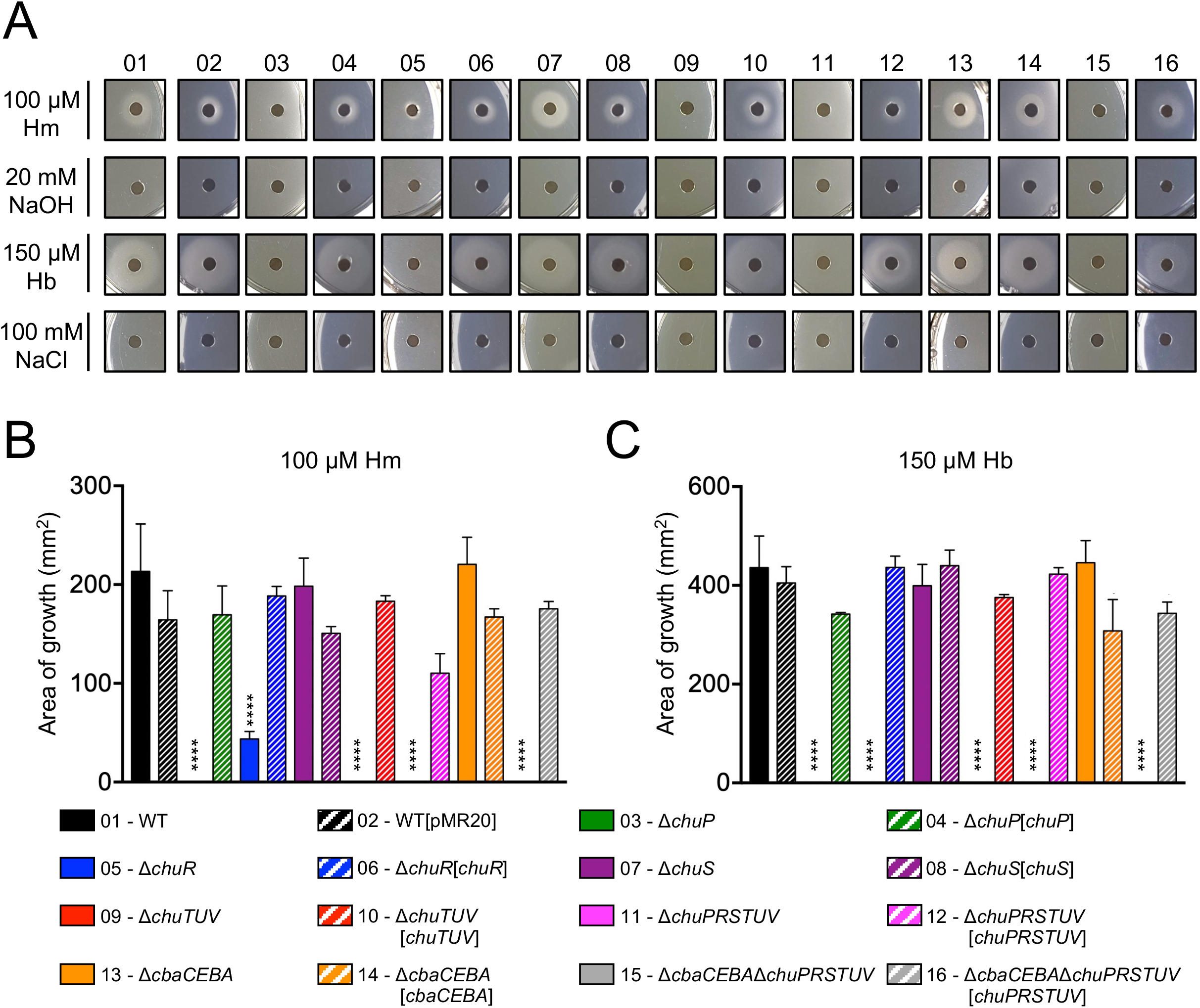
The *chu* operon encodes a heme uptake system (ChuRTUV) and a regulatory protein (ChuP) required for heme and hemoglobin utilization. **(A)** Nutrition assay for Hm and Hb under DP-imposed iron deficiency. The indicated strains were embedded into M9CH medium supplemented with 50 (Δ*cbaCEBA*), 100 (Δ*cbaCEBA*Δ*chuPRSTUV*), 125 (WT, Δ*chuP*, Δ*chuR*, Δ*chuS*, Δ*chuPRSTUV*, and all complemented strains) or 150 *μ*M DP (Δ*chuTUV*). Aliquots of 100 *μ*M Hm and 150 *μ*M Hb were provided as iron sources, while 20 mM NaOH and 100 mM NaCl were used as negative controls. Growth halos around the discs indicate compound utilization. Representative images are shown. **(B and C)** Quantification of Hm and Hb-stimulated growth. The area of the growth halos stimulated by Hm **(B)** and Hb **(C)** was measured using Image J software by subtracting the area of the discs. Data are from three biological replicates. Mutant and complemented strains were compared to WT and WT[pMR20], respectively. ****, p < 0.0001; when not shown, n.s. (not significant). One-way ANOVA followed by Tukey’s multiple-comparison test.

Considering that the Δ*chuS* strain showed no altered phenotype for Hm and Hb utilization (Figure 2), we evaluated its role on cell viability under heme excess (Supplementary Figure 2). However, growth defects were not observed for the WT, Δ*chuS*, and all mutant strains even at a high Hm concentration of 2 mM, indicating that the *chu* operon has no role during our heme excess conditions. Interestingly, deletion of the *chuPRSTUV* operon in the Δ*cbaCEBA* mutant strain improved the small colony size phenotype (Supplementary Figure 2), previously described for this strain (Batista et al., 2019). We also tested the hemolytic activity of the *chu* mutants on sheep-blood agar (Supplementary Figure 3). The strains Δ*chuR* (increased and intense halo) and Δ*cbaCEBA* (intense halo) showed altered hemolytic activity when compared to that of the WT and the other mutant strains. Although the meaning of these findings is unclear, we speculate that the increased hemolytic activity in these strains is a compensatory mechanism to deal with iron/heme scarcity.

### The Δ*chuP* mutant has altered activity of the siderophore viobactin

To verify whether the *chu* operon affects the siderophore activity in *C. violaceum*, we tested the *chu* mutants on PSA-CAS plates for siderophore detection as orange halos (Figure 3) as previously described (Batista et al., 2019). The Δ*chuP* and Δ*chuPRSTUV* mutants showed increased siderophore activity, while the Δ*chuR*, Δ*chuS*, and Δ*chuTUV* mutants had siderophore activity similar to that of the WT strain (Figure 3A, B). The Δ*cbaCEBA* strain showed no siderophore activity, as previously demonstrated (Batista et al., 2019), as well as the Δ*cbaCEBA*Δ*chuPRSTUV* strain (Figure 3A, B). After complementation, the siderophore activity was restored to WT levels in the Δ*chuP*[*chuP*] strain. For the strains Δ*chuPRSTUV*[*chuPRSTUV*] (almost absence of halo) and Δ*cbaCEBA*[*cbaCEBA*] (increased halo), the siderophore phenotypes reverted further on that observed in the WT strain (Figure 3A, B), perhaps owing to overexpression of the genes into the plasmid. These data indicate that the small regulatory protein ChuP controls siderophore activity in *C. violaceum*.

**Figure 3.**
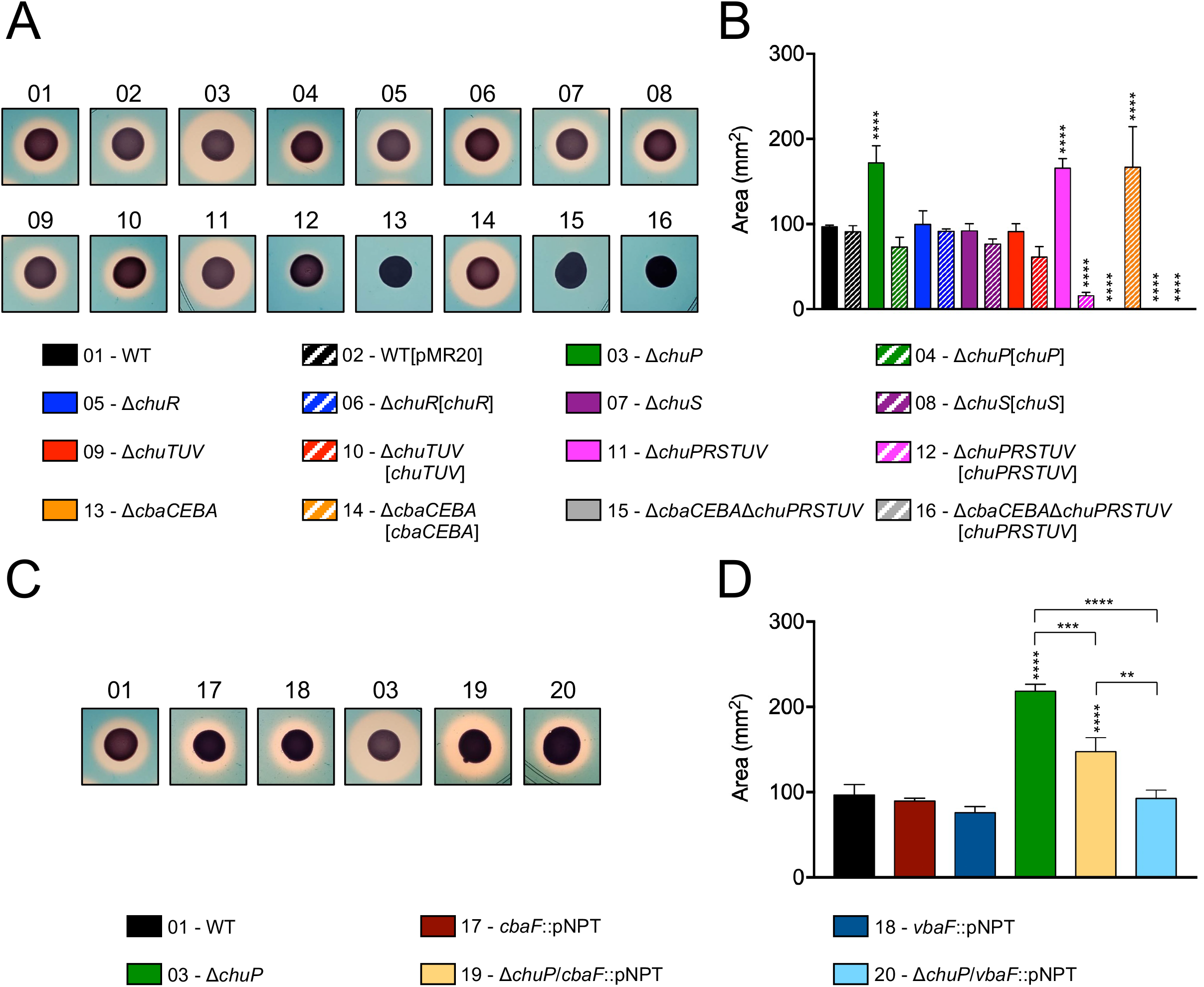
Deletion of *chuP* impacts siderophore activity in *C. violaceum*. **(A and B)** Role of the *chu* operon on siderophore activity in *C. violaceum*. Mutant strains without *chuP* showed increased siderophore halos. **(C and D)** The effect of ChuP occurs on the siderophore viobactin. For all indicated strains, the siderophore activity was determined by CAS assays on PSA-CAS plates. The orange halos indicating siderophore activity were photographed **(A and C)** and measured **(B and D)**, using Image J software. The area of the siderophore halos was calculated subtracting the area of bacterial growth. Data are from three biological replicates. Mutant and complemented strains **(B)** were compared to WT and WT[pMR20], respectively. Insertion mutants **(D)** were compared to the strains they derived from. **, p < 0.01; ***, p < 0.001; ****, p < 0.0001; when not shown, n.s. (not significant). Vertical asterisks indicate comparisons with the WT strain. One-way ANOVA followed by Tukey’s multiple-comparison test.

*C. violaceum* produces the catecholate-type siderophores chromobactin and viobactin employing the NRPS enzymes CbaF and VbaF, respectively (Batista et al., 2019). We combined mutation in *chuP* with mutations in *cbaF* or *vbaF* to understand which siderophore contributes to the increased siderophore activity in Δ*chuP*. Individual deletion of *cbaF* or *vbaF* genes in the WT had no effect on siderophore activity (Figure 3C, D), as previously reported (Batista et al., 2019). When these genes were deleted in the Δ*chuP* mutant background, a decrease in siderophore activity was observed in both cases. However, the halos were similar to that of the WT strain only when *vbaF* was deleted (Figure 3C, D), demonstrating a prominent role of viobactin on the increased siderophore activity of Δ*chuP*. Altogether, these results suggest that ChuP controls siderophore activity by acting over the synthesis and/or uptake of the siderophore viobactin.

### ChuP is a heme-binding post-transcriptional regulator of *chuR* and *vbuA* encoding TBDRs for heme and the siderophore viobactin

Our data indicate that mutation of *chuP* in *C. violaceum* abolished heme utilization (Figure 2) and altered activity of the siderophore viobactin (Figure 3). We employed different methodologies to elucidate how ChuP regulates these processes (Figure 4). First, we tested whether ChuP is a heme-binding protein. We purified the recombinant protein His-ChuP and performed a heme-binding assay (Figure 4A). After incubation with increasing Hm concentrations, a Soret peak at 413 nm was detected, indicating the formation of a ChuP-heme complex (Figure 4A). The differential absorption spectroscopy at 413 was used to fit a single binding model and determined that ChuP binds heme with a k_d_ = 18.36 ± 4.66 µM (Figure 4A, insert). Considering that HemP/HmuP proteins have been described as transcriptional activators (Escamila-Hernandez and O’Brian, 2012; Sato et al., 2017), we tested whether the *C. violaceum* ChuP regulates and binds into the intergenic regions upstream of *chuP* (promoter of the *chu* operon) and *chuR* (Figure 4B, C). The WT and Δ*chuP* strains harboring these constructs (p*chuP*-*lacZ* or p*chuR*-*lacZ*) were assessed by *β*-galactosidase assay under different iron levels. The p*chuP*-*lacZ* promoter fusion was highly active under iron deficiency (M9CH) with a gradual decrease upon Hm and FeSO_4_ supplementation (Figure 4B), as previously observed in the WT strain (Figure 1C). However, the same activity pattern was detected in the Δ*chuP* mutant strain, indicating that ChuP does not seem to regulate the promoter of the *chu* operon (Figure 4C). The p*chuR*-l*acZ* fusion had no promoter activity regardless of the strain or condition, indicating the absence of a promoter upstream of *chuR*. Therefore, this fusion is not useful to verify the effect of ChuP on *chuR* expression. Consistent with the *β*-galactosidase assays, our EMSA assays indicated that ChuP does not bind to the probes containing only the promoter of the *chu* operon or containing the entire region from *chuP* to *chuR* (Figure 4C). Altogether, these results demonstrate that ChuP does not regulate the promoter of the *chu* operon nor act as a DNA binding protein.

**Figure 4.**
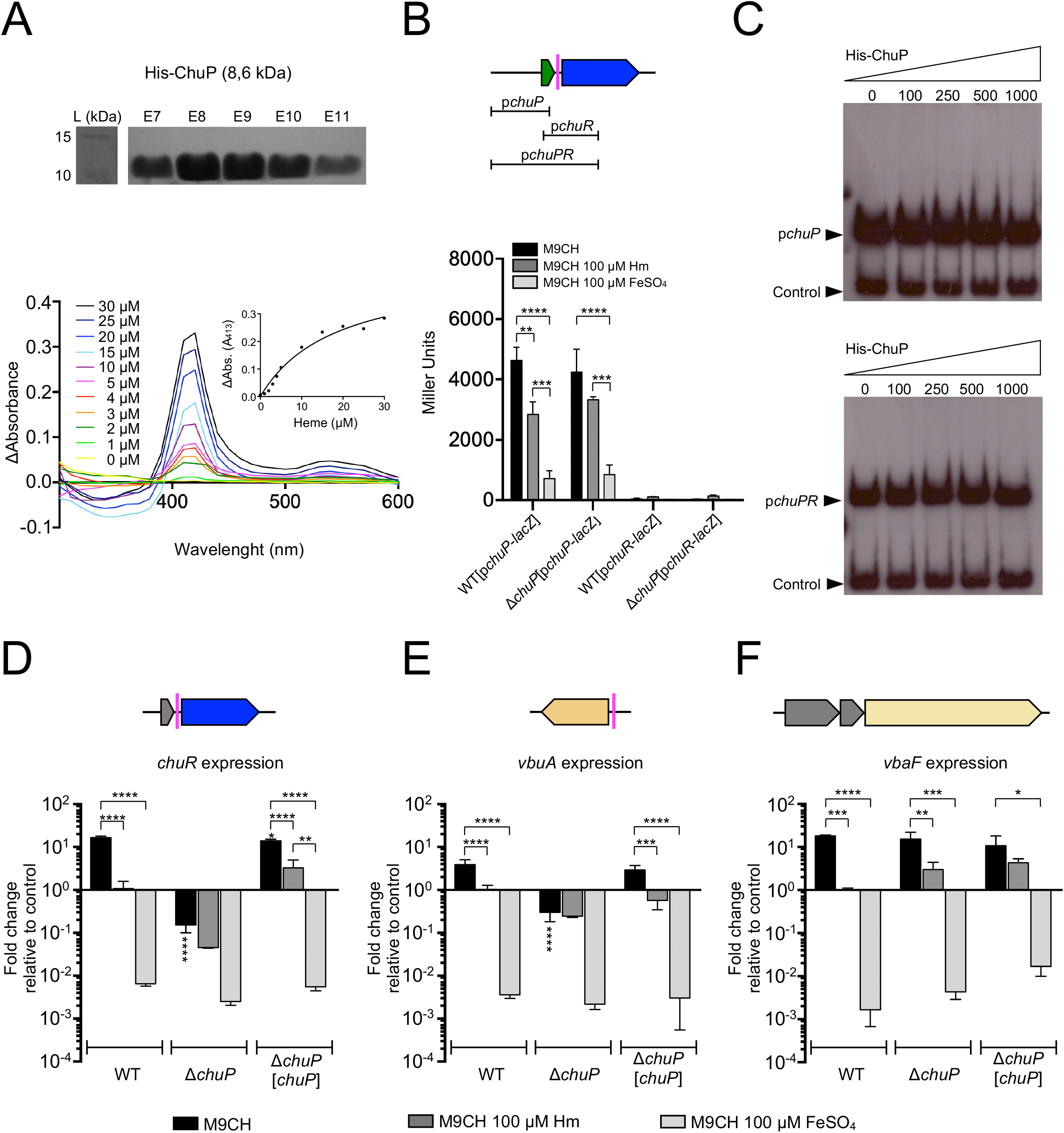
ChuP is a heme-binding regulatory protein that controls *chuR* and *vbuA* expression at a post-transcriptional level. **(A)** ChuP binds heme. The His-ChuP protein was purified (top) and incubated (10 *μ*M protein) with the indicated concentrations of Hm (bottom). The absorption spectra, recorded from 300 nm to 600 nm, revealed a peak at 413 nm, which was used to calculate the ChuP-Hm affinity (insert). Data are from a single experiment of three independent replicates. L, protein ladder; E7 to E11, eluted fractions of purified ChuP. **(B)** ChuP does not regulate the promoter of the *chu* operon. The scheme (top) indicates the regions used for *β*-galactosidase or EMSA assays. *β*-galactosidase assays were performed from the WT and Δ*chuP* strains harboring *chuP-lacZ* and *chuR*-*lacZ* fusions grown in M9CH in the indicated conditions of iron availability. Data are from three biological replicates. **, p < 0.01; ***, p < 0.001; ****, p < 0.0001. When not shown, n.s. (not significant). Two-way ANOVA followed by Dunnett’s multiple-comparison test. **(C)** ChuP does not bind to DNA probes covering from *chuP* to *chuR*. The indicated concentrations of His-ChuP were used in EMSA assays with the *chu* indicated probes. In both cases, the promoter region of CV_2599 (control) was used as an in-reaction unspecific negative control. **(D-F)** *chuR* and *vbuA* but not *vbaF* have HPRE sequences and are regulated by ChuP. The predicted HPREs are indicated as colored bars in the gene maps. Expression was evaluated by RT-qPCR. cDNA was reverse transcribed from RNA obtained from the WT, Δ*chuP*, and Δ*chuP*[*chuP*] strains grown in M9CH, and either untreated or treated with 100 *μ*M Hm or 100 *μ*M FeSO_4_. Expression of *chuR, vbuA*, and *vbaF* is shown as the fold change relative to the control condition (WT in M9CH 100 *μ*M Hm). Data are from three biological replicates. ****, p < 0.0001; *** p < 0.001, **, p < 0.01; *, p < 0.05; when not shown, n.s. (not significant). Vertical asterisks indicate comparisons with the WT strain at the same condition. One-way ANOVA followed by Tukey’s multiple-comparison test.

In *E. meliloti*, the HmuP protein activates the expression of the TBDR ShmR at a post-transcriptional level, probably by acting on a sequence HPRE (HmuP-responsive element). The HPRE sequences were predicted upstream of genes encoding heme TBDRs in many bacteria, including the *C. violaceum chuR* (sequence <u>CCCGCAAGCC</u>AGCCGACAG<u>CCAGCCAGCG</u>, -26 nt from the ATG start codon) (Amarelle et al., 2019). In addition to *chuR*, we found an HPRE sequence upstream of *vbuA* (sequence <u>GCCAGCC</u>AGACGAC<u>GCCGCCG</u>, -49 nt from the ATG start codon) (Figure 4D, E), raising the possibility that ChuP is a post-transcriptional regulator in *C. violaceum*. To verify this hypothesis, we performed RT-qPCR for *chuR, vbuA*, and *vbaF* genes with RNA harvested from the WT, Δ*chuP*, and Δ*chuP*[*chuP*] under different iron levels (Figure 4D, E, F). The expression of the three genes in the WT strain was high under iron-depleted and low under iron-sufficient conditions when compared to the control condition (WT grown in M9CH 100 µM Hm), as expected for genes related to iron acquisition (Figure 4D, E, F). Consistent with our phenotypic results and the presence of HPRE elements, the expression of *chuR* and *vbuA* was decreased in the Δ*chuP* strain regardless of the iron levels (Figure 4D, E), indicating that ChuP is a positive regulator required for the maximum expression of *chuR* and *vbuA* under iron limitation. Complementation of Δ*chuP* restored the expression of *chuR* and *vbuA* to the levels found in the WT strain (Figure 4D, E). No differences in *vbaF* expression were detected between the WT and Δ*chuP* strains in any of the tested conditions (Figure 4F), indicating that ChuP does not control the expression of VbaF, the NRPS for viobactin synthesis. Therefore, the decreased expression of *chuR* and *vbuA* in Δ*chuP* explains the inability of this mutant strain to use Hm and Hb (Figure 2) (via ChuR) and its increased siderophore halos (Figure 3) (inability to uptake viobactin via VbuA). Altogether, these results demonstrate that ChuP integrates the acquisition of heme and siderophore by acting as a heme-binding post-transcriptional regulator of the TBDR genes *chuR* and *vbuA*.

### *C. violaceum* employs both siderophores and heme for iron acquisition during infection

To assess the role of the heme utilization system ChuPRSTUV during *C. violaceum* infection, we performed mice virulence assays (Figure 5). The animals were i.p. injected with a dose of 10^6^ bacterial cells and analyzed for survival during seven days post-infection (Figure 5A, B). The five null mutant strains of the ChuPRSTUV system showed barely or no virulence attenuation compared to the *C. violaceum* WT strain (Figure 5A). Previously, we determined that abrogating siderophore production in *C. violaceum* by deletion of the *cbaCEBA* genes causes moderate attenuation in virulence (Batista et al., 2019). Therefore, we checked whether heme and siderophores cooperate for virulence. Indeed, the Δ*cbaCEBA* strain showed an intermediate virulence attenuation, as expected, while a more expressive virulence attenuation was observed for the Δ*cbaCEBA*Δ*chuPRSTUV* strain (Figure 5A). Complementation of the latter strain with the *chuPRSTUV* operon reverted its virulence attenuation phenotype to the pattern observed for the Δ*cbaCEBA* mutant (Figure 5B).

**Figure 5.**
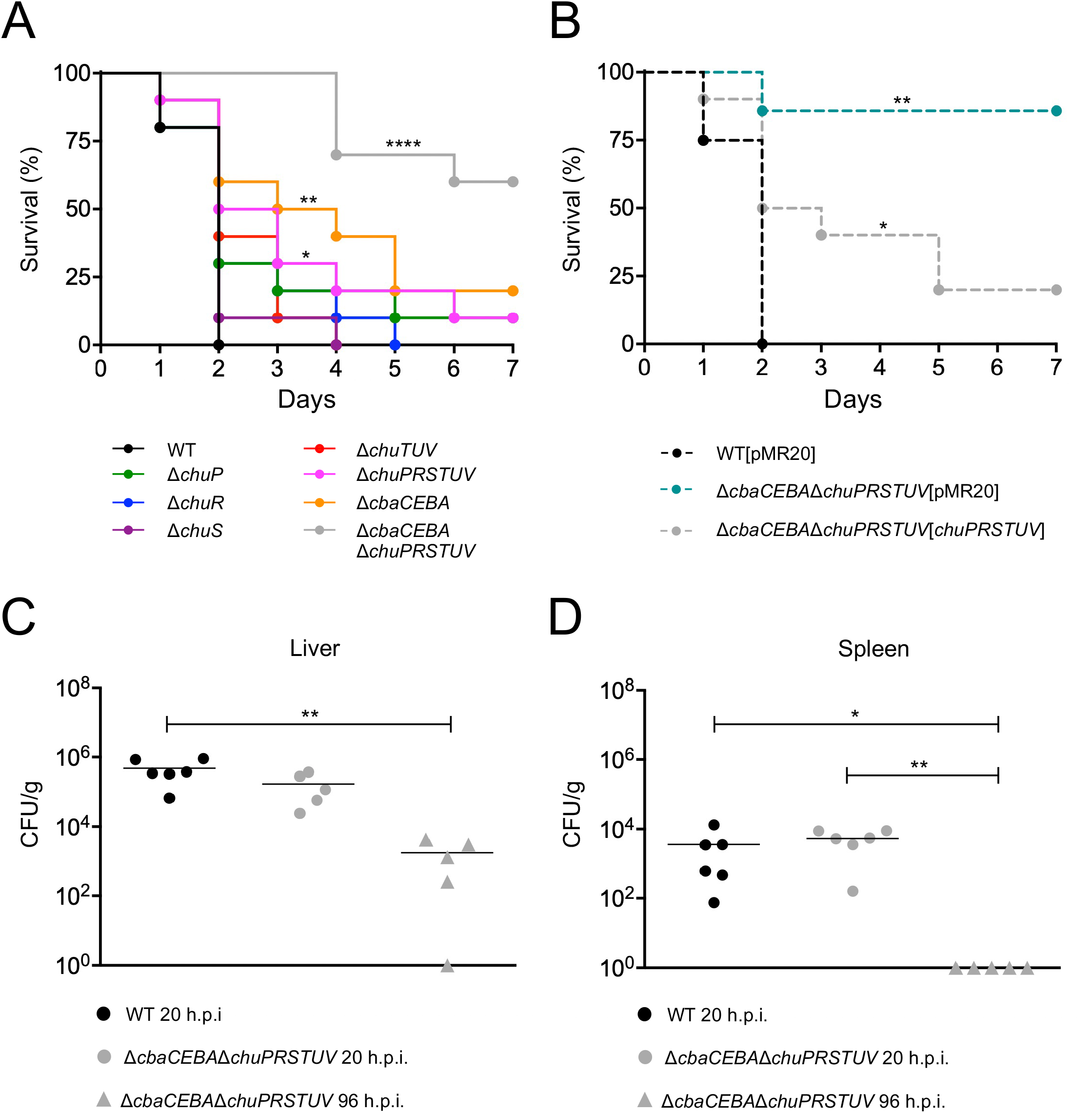
*C. violaceum* requires siderophores and heme but prefers siderophores in a mice model of acute infection. (**A and B)** Survival curves of infected BALB/c mice. Animals (n = 8 for WT[pMR20]; n = 7 for Δ*cbaCEBA*Δ*chuPSRTUV*[*chuPRSTUV*]; n = 10 for all other strains) were i.p. injected with 10^6^ CFU of the indicated *C. violaceum* mutant **(A)** and complemented **(B)** strains. Animal survival was monitored daily for a week. *, p < 0.05; **, p < 0.01; ****, p < 0.0001; when not shown, n.s. (not significant). Log-rank (Mantel-Cox) test. **(C and D)** Bacterial burden in organs. Animals were infected with 10^6^ CFU of the indicated strains. After 20h or 96 h post-infection (h.p.i.), the liver **(C)** and the spleen **(D)** were collected, homogenized, serially diluted, and plated for CFU quantification. *, p < 0.05; **, p < 0.01; when not indicated, n.s. (not significant). One-way ANOVA followed by Tukey’s (liver) or Dunnett’s (spleen) multiple-comparison tests.

We evaluated the bacterial burden in the liver and spleen, two organs colonized during *C. violaceum* infection that are involved in host heme recycling. Interestingly, the Δ*cbaCEBA*Δ*chuPRSTUV* mutant displayed the same CFU counting as the WT strain at 20 hours post-infection in both organs (Figure 5C, D). However, the bacterial burden was reduced (in the liver) and eliminated (in the spleen) at 96 hours post-infection (Figure 5C, D). These results indicate that the absence of siderophores and heme uptake does not impair initial colonization but impairs the bacterial maintenance in later infection stages. Altogether, these results indicate an interplay between the iron-acquisition strategies based on siderophore and heme during *C. violaceum* infection. In our acute infection model, the requirement of heme uptake for virulence becomes evident in the absence of siderophores.

## DISCUSSION

In this work, we identified and characterized a heme and hemoglobin utilization system, here named ChuPRSTUV, which connects, via the regulatory protein ChuP, iron acquisition by heme and siderophore during *C. violaceum* infection (Figure 6). Our data indicated that the genes *chuPRSTUV* compose an operon repressed by Fur under iron sufficiency. During iron limitation (as found inside the host), high expression of the *chu* operon (for heme uptake by the transport system ChuR-ChuTUV) and the *vbaF* and *vbuA* genes (for synthesis and uptake of the siderophore viobactin) occurred (Figure 6A, B). Remarkably, we found that the maximum expression in iron scarcity of the TBDR genes *chuR and vbuA* depends on the small heme-binding protein ChuP. In our model, we propose that ChuP is a positive post-transcriptional regulator acting in the 5’-UTR of the *chuR and vbuA* transcripts (Figure 6A). Without ChuP, the expression of *chuR* and *vbuA* dropped, rendering the Δ*chuP* mutant strain its inability to use Hm and Hb via ChuR and its increased siderophore halos (deficiency to uptake viobactin via VbuA). Moreover, our virulence data in mice demonstrated that *C. violaceum* uses both heme and siderophore for iron acquisition during infection, with a preference for siderophores over the Chu heme uptake system (Figure 6C).

**Figure 6.**
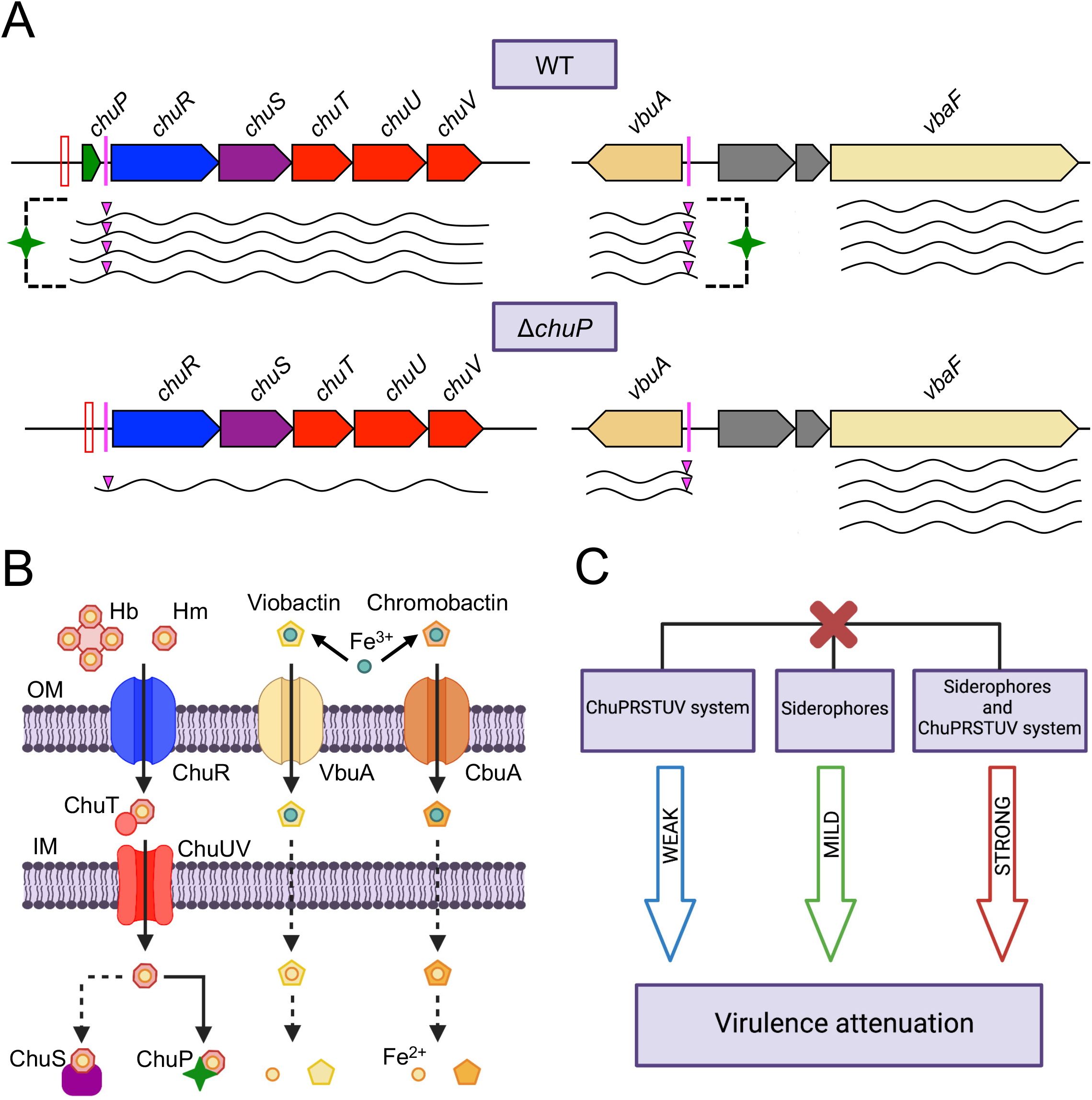
Model of how *C. violaceum* connects iron acquisition by heme and siderophore during infection. **(A)** Regulatory function of ChuP over *chuR* and *vbuA*. **(B)** Iron acquisition systems in *C. violaceum* for the uptake of heme (this work) and the siderophores viobactin and chromobactin (Batista et al., 2019). **(C)** Interplay between the uptake of heme and siderophore in the *C. violaceum* virulence. We propose that ChuP links heme and siderophore utilization by acting as a positive regulator of *chuR* and *vbuA*, which encode TBDRs for the uptake of heme (ChuR) and the siderophore viobactin (VbuA). In addition to Fur derepression, the expression of *chuR* and *vbuA* depends on ChuP under iron deficiency, possibly by a post-transcriptional mechanism involving HPRE (HmuP-responsive elements) sequences found in the 5’-UTR of the *chuR* and *vbuA* transcripts. In the absence of ChuP, the abundance of *chuR* and *vbuA* transcripts decreases, causing a reduction in heme and viobactin utilization. Different levels of virulence attenuation occurred when the ChuPRSTUV system (weak), the siderophores (mild), or both (strong) were deleted, indicating that *C. violaceum* prefers siderophores over heme during infection but relies on heme in the absence of siderophores. Red bars, Fur boxes; Pink bars and triangles, predicted HPRE sequences; Dashed lines, unknown mechanisms.

We demonstrated that the *chuPRSTUV* genes are co-transcribed from an iron-responsive and Fur-repressed promoter in *C. violaceum*, a gene cluster organization and expression pattern that fit with those found for heme uptake system in other bacteria, such as *B. multivorans, Yersinia* spp., and *P. aeruginosa* (Sato et al., 2017; Si et al., 2017; Schwisow et al., 2018; Otero-Asman et al., 2019). Our nutrition assays indicated that, with the exception of *chuS*, all genes of the *chu* operon are required for heme and hemoglobin utilization in *C. violaceum*, suggesting that ChuRTUV, composed by the TBDR ChuR and the ABC transport system ChuTUV, is a heme uptake system. This mechanism of heme import across the cell envelope is found in many Gram-negative bacteria (Stojiljkovic and Hantke, 1992; Burkhard et al., 2007; Balhesteros et al., 2017; Huang and Wilks, 2017). The mutant Δ*chuR* but not the mutant Δ*chuTUV* showed a small halo of heme utilization, and both mutants were unable to use hemoglobin, suggesting that *C. violaceum* maybe have another TBDR for heme uptake but relies specifically on ChuR to obtain heme from hemoglobin. Indeed, in *P. aeruginosa*, a bacterium with three heme uptake systems (Phu, Has, and Hxu), the same ABC-transport system (PhuTUV) transfers heme to the cytosol after uptake by the TBDRs PhuR and HasR (Ochsner et al., 2000; Smith and Wilks, 2015; Otero-Asman et al., 2019). ChuR appears to act as a direct heme uptake transporter given that we do not find genes encoding hemophores in *C. violaceum* and ChuR does not have an N-terminal extension typically found in hemophore-based heme uptake systems (Biville et al., 2004; Wandersman and Delapelaire, 2012; Huang and Wilk, 2017). Since the *C. violaceum* Δ*chuS* mutant showed no phenotype under heme limitation or excess, further biochemical studies are necessary to investigate whether ChuS is a heme chaperone or a non-canonical heme oxygenase involved in heme degradation, as described in other bacteria (Suits et al., 2005; Amarelle et al., 2016; Lee et al., 2017).

Recent studies have shown regulatory and functional connections between heme and siderophores (Otero-Asman et al., 2019; Zygiel et al., 2020; Batko et al., 2021; Glanville et al., 2021). The increased siderophore halos detected in Δ*chuP* and Δ*chuPRSTUV* mutant strains indicate that *chuP* is the gene of the *chu* operon that connects heme and siderophore utilization in *C. violaceum*. ChuP belongs to the HemP/HmuP protein family, whose members are found in many proteobacteria (Amarelle et al., 2019). However, only HmuP from *S. meliloti* and *B. japonicum* and HemP from *B. multivorans* have been characterized. They are small regulatory proteins required for heme utilization by acting as positive regulators of heme-acquisition TBDR genes (Amarelle et al., 2010; Escamila-Hernandez and O’Brian, 2012; Sato et al., 2017; Amarelle et al., 2019). In *B. multivorans*, a *hmuP* mutant showed decreased siderophore halos, but the underlying mechanism remains unexplored (Sato et al., 2017). Our data indicate that ChuP links heme and siderophore utilization by acting as a positive regulator required for the expression of *chuR* and *vbuA*, genes encoding the TBDRs used by *C. violaceum* for the uptake of heme/hemoglobin (ChuR) and the siderophore viobactin (VbuA) (Batista et al., 2019). Our data favor a working model of ChuP as a heme-binding post-transcriptional regulator acting in the 5’-UTR of the *chuR and vbuA* transcripts (Figure 6). Supporting this model, we found that (i) ChuP of *C. violaceum* binds heme, as demonstrated for HmuP of *B. multivorans* (Sato et al., 2017); (ii) ChuP does not regulate the promoter of the *chu* operon (in front of *chuP*) and its effect on *chuR* does not occur at the transcriptional level since there is no promoter in front of *chuR* and ChuP does not bind to DNA probes covering the entire region from *chuP* to *chuR*; (iii) there is the presence, upstream of *chuR* and *vbuA*, of HPRE elements, which were described as conserved sequences probably acting on mRNA in the 5’-UTR of genes encoding heme-related TBDRs (Amarelle at al., 2019). Although HemP/HmuP proteins lack a typical DNA binding domain, they were described as direct DNA binding proteins in *B. japonicum* and *B. multivorans*, maybe by interacting with Irr and Fur (Escamilla-Hernandez and O’Brian, 2012; Sato et al., 2017). Our results suggest that ChuP in *C. violaceum* works similarly to HmuP in *E. meliloti*, but more work is necessary to understand how ChuP exerts its role as a post-transcriptional regulator on its target genes.

Several investigations have found that genes encoding heme uptake systems are upregulated *in vivo* (Cook et al., 2019; Rivera-Chávez and Mekalanos, 2019) and required for colonization and virulence of many bacterial pathogens (Skaar et al., 2004; Si et al., 2017; Abdelhamed et al., 2018; Cook et al., 2019; Rivera-Chávez and Mekalanos, 2019; Chatterjee et al., 2020). In many cases, bacteria explore multiple host iron sources, employing both heme and siderophore-based iron acquisition systems (Contreras et al., 2014; Huang and Wilks, 2017). Our prior work revealed that *C. violaceum* requires catecholate-type siderophores for virulence in mice (Batista et al., 2019). Our current findings based on the characterization of mutants without either siderophores, the *chu* operon, or both indicate that *C. violaceum* uses siderophores and heme but prioritizes siderophores over heme as an iron source during infection, at least in our mice model of acute systemic infection (Figure 6C). In agreement with our data, a study that characterized mutants of multiple iron uptake systems showed a clear predominance of siderophores over heme transport systems in *P. aeruginosa* infecting lung (Minandri et al., 2016). However, the preference for a particular iron source changes according to its availability or the infection context. For instance, *S. aureus* prefers heme but uses siderophores when heme is scarce (Skaar et al., 2004); *P. aeruginosa* prioritizes siderophore systems in acute infections but switches to heme in long-term chronic infections (Marvig et al., 2014; Nguyen et al., 2014); and *Vibrio cholerae* relies on heme released by cholera toxin-dependent damage in the intestine (Rivera-Chávez and Mekalanos, 2019). Currently, we are developing a mouse model of abscess for *C. violaceum* infection. It will be interesting to investigate in this model whether *C. violaceum* alters its preference for siderophores and heme in log-term infections.

## ACKNOWLEDGMENTS

This research was supported by grants from the São Paulo Research Foundation (FAPESP; grants 2018/01388-6 and 2020/00259-8) and Fundação de Apoio ao Ensino, Pesquisa e Assistência do Hospital das Clínicas da FMRP-USP (FAEPA). During this work VML (2018/17716-2) and BBB (2018/19058-2) were supported by FAPESP fellowships.

## AUTHOR CONTRIBUTIONS

JFSN and VML conceived and designed the study. VML and BBB performed the experiments. VML and JFSN performed data analysis and interpretation. VML and JFSN wrote the paper.

## REFERENCES

Abdelhamed, H., Ibrahim, I., Baumgartner, W., Lawrence, M.L., Karsi, A. (2018). The virulence and immune protection of Edwardsiella ictaluri HemR mutants in catfish. Fish Shellfish Immunol. 72, 153–160. doi: 10.1016/j.fsi.2017.10.041.

Amarelle, V., Koziol, U., Rosconi, F., Noya, F., O’Brian, M.R., Fabiano, E. (2010). A new small regulatory protein, HmuP, modulates haemin acquisition in Sinorhizobium melioti. Microbiology 156, 1873–1882. doi: 10.1099/mic.0.037713-0.

Amarelle, V., Rosconi, F., Lázaro-Martinez, J.M., Buldain, G., Noya, F., O’Brian, M.R. et al. (2016). HmuS and HmuQ of Ensifer/Sinorhizobium meliloti degrade heme in vitro and participate in heme metabolism in vivo. Biometals 29, 333–347. doi: 10.1007/s10534-016-9919-3

Amarelle, V., Koziol, U., Fabiano, E. (2019). Highly conserved nucleotide motifs present in the 5’UTR of the heme-receptor gene shmR are required for HmuP-dependent expression of shmR in Ensifer meliloti. Biometals 32, 273–291. doi: 10.1007/s10534-019-00184-6.

Balhesteros, H., Shipelskiy, Y., Long, N. J., Majumdar, A., Katz, B.B., Santos, N.M. et al. (2017). TonB-dependent heme/hemoglobin utilization in Caulobacter crescentus HutA. J. Bacteriol. 199:e00723–16. doi: 10.1128/JB.00723-16.

Batista, B.B., Santos, R.E.R.S., Ricci-Azevedo, R., da Silva Neto, J.F. (2019). Production and uptake of distinct endogenous cathecolate-type siderophores are required for iron acquisition and virulence in Chromobacterium violaceum. Infect. Immun. 87:e00577–19. doi: 10.1128/IAI.00577-19.

Batista, J.H., da Silva Neto, J.F. (2017). Chromobacterium violaceum pathogenicity: updates and insights from genome sequencing of novel Chromobacterium species. Front. Micriobiol. 8:2213. doi: 10.3389/fmicb.2017.02213.

Batista, J.H., Leal, F.C., Fukuda, T.T., Diniz, J.A., Almeida, F., Pupo, M.T. et al. (2020). Interplay between two quorum sensing-regulated pathways, violacein biosynthesis and VacJ/Yrb, dictates outer membrane vesicles biogenesis in Chromobacterium violaceum. Envrionm. Micriobiol. 22, 2432–2442. doi: 10.1111/1462-2920.15033.

Batko I.Z., Flannagan, R.S., Guariglia-Oropeza, V., Sheldon, J.R., Heinrichs, D.E. (2021). Heme-ependent siderophore utilization promotes iron-restricted growth of the Staphylococcus aureus hemB small-colony variant. J Bacteriol. 203:e0045821. doi:10.1128/JB.00458-21

Biville, F., Cwerman, H., Létoffé, S., Rossi, M.S., Drouet, V., Ghigo, J.M. et al. (2004). Haemophore-mediated signaling in Serratia marcescens: a new mode of regulation for an extra cytoplasmic function (ECF) sigma factor involved in haem acquisition. Mol. Microbiol. 53, 1267–1277. doi: 10.1111/j.1365-2958.2004.04207.x.

Braun, V., Hantke, K. (2011). Recent insights into iron import by bacteria. Curr. Opin. Chem. Biol. 15, 328–334. doi: 10.1016/j.cbpa.2011.01.005.

Brazilian National Genome Project Consortium. (2003). The complete genome sequence of Chromobacterium violaceum reveals remarkable and exploitable bacterial adaptability. Proc. Natl. Acad. Sci. USA. 100, 11660–11665. doi: 10.1098/rspb.2013.1055.

Burkhard, K.A., Wilks, A. (2007). Characterization of the outer membrane receptor ShuA from the heme uptake system of Shigella dysenteriae. Substrate specificity and identification of the heme protein ligands. J. Biol. Chem. 18, 15126–15136. doi: 10.1074/jbc.M611121200.

Cassat, J.E., Skaar, E.P. (2013). Iron in Infection and Immunity. Cell Host & Microbe 13, 509–519. doi: 10.1016/j.chom.2013.04.010.

Chandrangsu P., Rensing, C., Helmann, J.D. (2017) Metal homeostasis and resistance in bacteria. Nat. Rev. Microbiol. 15: 338–350. doi: 10.1038/nrmicro.2017.15.

Chatterjee, N., Cook, L.C.C., Lyles, K.V., Nguyen, H.A., Devlin, D.J., Thomas, L.S. et al. (2020). A novel heme transporter from the energy coupling factor family is vital for group A Streptococcus colonization and infections. J Bacteriol. 202:e00205–20. doi:10.1128/JB.00205-20

Choby, J.E., Skaar, E.P. (2016). Heme synthesis and acquisition in bacterial pathogens. J. Mol. Biol. 428, 3408–3428. doi: 10.1016/j.jmb.2016.03.018.

Contreras, H., Chim, N., Credali, A., Goulding, C.W. (2014). Heme uptake in bacterial pathogens. Curr. Opin. Chem. Biol. 19, 34–41. doi: 10.1016/j.cbpa.2013.12.014.

Cook, L.C.C., Chatterjee, N., Li, Y., Andrade, J., Federle, M.J., Eichenbaum, Z. (2019). Transcriptomic analysis of Streptococcus pyogenes colonizing the vaginal mucosa identifies hupY, an MtsR-regulated adhesin involved in heme utilization. mBio. 10:e00848–19. doi:10.1128/mBio.00848-19

da Silva Neto, J.F., Braz, V.S., Italiani, V.C.S., Marques, M.V. (2009). Fur controls iron homeostasis and oxidative stress defense in the oligotrophic alpha-proteobacterium Caulobacter crescentus. Nucleic Acids Res. 37, 4812–4825. doi: 10.1093/nar/gkp509.

da Silva Neto, J.F., Negretto, C.C., Netto, L.E.S. (2012). Analysis of the organic hydroperoxide response of Chromobacterium violaceum reveals that OhrR is a Cys-based redox sensor regulated by thioredoxin. PLoS One 7:e47090. doi: 10.1371/journal.pone.0047090.

Eakanunkul, S., Lukat-Rodgers, G.S., Sumithran, S., Ghosh, A., Rodgers, K.R., Dawson, J.H. et al. (2005). Characterization of the periplasmic heme-binding protein ShuT from the heme uptake system of Shigella dysenteriae. Biochem. 44, 13179–13191. doi: 10.1021/bi050422r.

Escamilla-Hernandez, R., O’ Brian, M.R. (2012). HmuP is a coactivator of Irr-dependent expression of heme utilization genes in Bradyrhizobium japonicum. J. Bacteriol. 194, 3137–3143. doi: 10.1128/JB.00071-12.

Fournier, C., Smith, A., Delepelaire, P. (2011). Haem release from haemopexin by HxuA allows Haemophilus influenzae to escape host nutritional immunity. Mol. Micribiol. 80, 133–148. doi: 10.1111/j.1365-2958.2011.07562.x.

Ganz, T., Nemeth, E. (2015). Iron homeostasis in host defence and inflammation. Nat. Rev. Immunol. 15, 500–510. doi: 10.1038/nri3863.

Glanville, D.G., Mullineaux-Sanders, C., Corcoran, C.J., Burger, B.T., Imam, S., Donohue, T.J. et al. (2021). A high-throughput method for identifying novel genes that influence metabolic pathways reveals new iron and heme regulation in Pseudomonas aeruginosa. mSystems. 6:e00933–20. doi: 10.1128/mSystems.00933-20.

Gober, J.W., Shapiro, L. (1992). A developmentally regulated Caulobacter flagellar promoter is activated by 3’ enhancer and IHF binding elements. Mol. Biol. Cell. 3, 913–926. doi: 10.1091/mbc.3.8.913.

Hanahan, D. (1983). Studies of transformation of Escherichia coli with plasmids. J. Mol. Biol. 166, 557–580. doi: 10.1016/S0022-2836(83)80284-8.

Hood, M.I., Skaar, E.P. (2012). Nutritional immunity: transition metals at the pathogen-host interface. Nat. Rev. Micribiol. 10, 525–537. doi: 10.1038/nrmicro2836.

Huang, W., Wilks, A. (2017). Extracellular heme uptake and the challenge of bacterial cell membranes. Annu. Rev. Biochem. 86, 799–823. doi: 10.1146/annurev-biochem-060815-014214.

Khalifa, S.M.A., Khaldi, T.A., Alqahtani, M.M., et al. (2015). Two siblings with fatal Chromobacterium violaceum sepsis linked to drinking water. BMJ Case Rep. 2015:bcr2015210987. doi: 10.1136/bcr-2015-210987.

Klebba, P.E., Newton, S.M.C., Six, D.A., Kumar, A., Yang, T., Nairn, B.L. et al. (2021). Iron acquisition systems of Gram-negative bacterial pathogens define TonB-dependent pathways to novel antibiotics. Chem Rev. 121, 5193–5239. doi: 10.1021/acs.chemrev.0c01005.

Kumar, M.R. (2012). Chromobacterium violaceum: a rare bacterium isolated from a wound over the scalp. Int. J. Appl. Basic Med. Res. 2, 70–72. doi: 10.4103/2229-516X.96814.

Lamattina, J.W., Nix, D.B., Lanzilotta, W.N. (2016). Radical new paradigm for heme degradation in Escherichia coli O157:H7. Proc. Natl. Acad. Sci. 113, 12138–12143. doi: 10.1073/pnas.1603209113.

Lee, M.J.Y., Wang, Y., Jiang, Y. Li, X., Ma, J., Tan, H. et al. (2017). Function coupling mechanism of PhuS and HemO in heme degradation. Sci. Rep. 7:1123. doi: 10.1038/s41598-017-11907-5.

Livak, K.J., Schmittgen, T.D. (2001). Analysis of relative gene expression data using real-time quantitative PCR and the 2(-Delta Delta C(T)) method. Methods. 25, 402–408. doi: 10.1006/meth.2001.1262.

Maltez, V.I., Tubbs, A.L., Cook, K.D., Aachoui, Y., Falcone, E.L., Holland, S.M. et al. (2015). Inflammasomes coordinate pyroptosis and natural killer cell cytotoxicity to clear infection by a ubiquitous environmental bacterium. Immunity 43, 987–997. doi: 10.1016/j.immuni.2015.10.010.

Marvig, R.L., Damkiær, S., Khademi, S.M., Markussen, T.M., Molin, S., Jelsbak, L. (2014). Within-host evolution of Pseudomonas aeruginosa reveals adaptation toward iron acquisition from hemoglobin. mBio. 5:e00966–14. doi:10.1128/mBio.00966-14.

Miki, T., Iguchi, M., Akiba, K., Hosono, M., Sobue, T., Danbara, H. et al. (2010). Chromobacterium pathogenicity island 1 type III secretion system is a major virulence determinant for Chromobacterium violaceum-induced cell death in hepatocytes. Mol. Microbiol. 77, 855–872. doi: 10.1111/j.1365-2958.2010.07248.x.

Minandri, F., Imperi, F., Frangipani, E., Bonchi, C., Visaggio, D., Facchini, M. et al. (2016). Role of iron uptake systems in Pseudomonas aeruginosa virulence and airway infection. Infect. Immun. 84, 2324–2335. doi: 10.1128/IAI.00098-16.

Nguyen, A.T., O’Neill, M.J., Watts, A.M., Robson, C.L., Lamont, I.L., Wilks, A. et al. (2014) Adaptation of iron homeostasis pathways by a Pseudomonas aeruginosa pyoverdine mutant in the cystic fibrosis lung. J Bacteriol. 196, 2265–2276. doi:10.1128/JB.01491-14

Noinaj, N., Guillier, M., Barnard, T. J., Buchanan, S.K. (2010). TonB-dependent transporters: regulation, structure and function. Ann. Rev. Microbiol. 64, 43–60. doi: 10.1146/annurev.micro.112408.134247.

Ochsner, U.A., Johnson, Z., Vasil, M.L. (2000). Genetics and regulation of two distinct haem-uptake systems, phu and has, in Pseudomonas aeruginosa. Microbiology 146, 185–198. doi: 10.1099/00221287-146-1-185.

Otero-Asman, J.R., García-García, A.I., Civantos, C., Quesada, J.M., Llamas, M.A. (2019). Pseudomonas aeruginosa possesses three distinct systems for sensing and using the host molecule haem. Environ. Microbiol. 12, 4629–4647. doi: 10.1111/1462-2920.14773.

Palmer, L. D., Skaar, E. P. (2016). Transition metals and virulence in bacteria. Annu. Rev. Genet. 50, 67–91. doi: 10.1146/annurev-genet-120215-035146.

Parrow, N. L., Flemming, R.E., Minnick, M.F. (2013). Sequestration and scavenging of iron in infection. Infect. Immun. 81, 3503–3514. doi: 10.1128/IAI.00602-13.

Previato-Mello, M., Meireles, D.A., Netto, L.E.S., da Silva Neto, J.F. (2017). Global transcriptional response to organic hydroperoxide and the role of OhrR in the control of virulence traits in Chromobacterium violaceum. Infect. Immun. 85:e00017–17. doi: 10.1128/IAI.00017-17.

Puri, S., O’Brian, M.R. (2006). The hmuQ and hmuD genes from Bradyrhizobium japonicum encode heme-degrading enzymes. J. Bacteriol. 188, 6476–6482. doi: 10.1128/JB.00737-06.

Rivera-Chávez, F., Mekalanos, J.J. (2019). Cholera toxin promotes pathogen acquisition of host-derived nutrients. Nature. 572, 244–248. doi:10.1038/s41586-019-1453-3

Roberts, R.C., Toochinda, C., Acedissian, M., Baldini, R.L., Gomes, S.L., Shapiro, L. (1996). Identification of a Caulobacter crescentus operon encoding hrcA, involved in negatively regulating heat-inducible transcription, and the chaperone gene grpE. J. Bacteriol. 178, 1829–1841. doi: 10.1128/jb.178.7.1829-1841.1996.

Runyen-Janecky, L.J. (2013). Role and regulation of heme iron acquisition in Gram-negative pathogens. Front. Cell. Infect. Microbiol. 3:55. doi: 10.3389/fcimb.2013.00055.

Santos, R.E.R.S., Batista, B.B., da Silva Neto, J. F. (2020). Ferric uptake regulator Fur coordinates siderophore production and defense against iron toxicity and oxidative stress and contributes to virulence in Chromobacterium violaceum. Applied Environm. Microbiol. 86:e01620–20. doi: 10.1128/AEM.01620-20.

Sarvan, S., Butcher, J., Stintzi, A., Couture, J.F. (2018). Variation on a theme: investigating the structural repertoires used by ferric uptake regulator to control gene expression. Biometals 31, 681–704. doi: 10.1007/s10534-018-0120-8.

Sato, T., Nonoyama, S., Kimura, A., Nagata, Y., Ohtsubo, Y., Tsuda, M. (2017). The small protein HemP is a transcriptional activator of the hemin uptake operon in Burkholderia multivorans ATCC 17616. App. Environm. Microbiol. 83:e00479–17. doi: 10.1128/AEM.00479-17.

Schwiesow, L., Metteret, E., Wei, Y., Miller, H.K., Herrera, N.G., Balderas, D. et al. (2018). Control of hmu heme uptake genes in Yersinia pseudotuberculosis in response to iron sources. Front. Cell. Infect. Microbiol. 8:47. doi: 10.3389/fcimb.2018.00047.

Sheldon, J.R., Laakso, H.A., Heinrichs, D.E. (2016). Iron acquisition strategies of bacterial pathogens. Microbiol. Spectr. 4:2. doi: 10.1128/microbiolspec.VMBF-0010-2015.

Si, M., Wang, Y., Zhang, B., Zhao, C., Kang, Y., Bai, H. et al. (2017). The type VI secretion system engages a redox-regulated dual-function heme transporter for zinc acquisition. Cell Rep. 20, 949–959. doi: 10.1016/j.celrep.2017.06.081.

Simon, R., Priefer, U., Pühler, A. (1983). A broad host range mobilization system for in vivo genetic engineering: transposon mutagenesis in Gram-negative bacteria. Nat. Biotechnol. 1, 784–791. doi: 10.1038/nbt1183-784.

Skaar, E.P. (2010). The battle for iron between bacterial pathogens and their vertebrate hosts. PLoS Pathog. 6:e1000949. doi: 10.1371/journal.ppat.1000949.

Skaar, E.P., Humayun, M., Bae, T., Debord, K. L., Schneewind, O. (2004). Iron-source preference of Staphylococcus aureus infections. Science 305, 1626–1628. doi: 10.1126/science.1099930.

Smith, A.D., Wilks, A. (2015). Differential contributions of the outer membrane receptors PhuR and HasR to heme acquisition in Pseudomonas aeruginosa. J. Biol. Chem. 290, 7756–7766. doi: 10.1074/jbc.M114.633495.

Stojiljkovic, I., Hantke, K. (1992). Hemin uptake system of Yersinia enterocolitica: similarities with other TonB-dependent systems in Gram-negative bacteria. EMBO Journal 11, 4359–4367. PMCID: PMC557009.

Suits, M.D., Pal, G.P., Nakatsu, K., Matte, A., Cygler, M., Jia, Z. (2005). Identification of an Escherichia coli O157:H7 heme oxygenase with tandem functional repeats. Proc. Natl. Acad. Sci. USA. 102, 16955–16960. doi: 10.1073/pnas.0504289102.

Wandersman, C., Delepelaire, P. (2012). Haemophore functions revisited. Mol. Microbiol. 85, 618–631. doi: 10.1111/j.1365-2958.2012.08136.x.

Yang, C.H., Li, Y.H. (2011). Chromobacterium violaceum infection: a clinical review of an important but neglected infection. J. Clin. Med. Assoc. 74, 435–441. doi: 10.1016/j.jcma.2011.08.013.

Zhao, Y., Yang, J., Shi, J., Gong, Y.N., Lu, Q., Xu, H. et al. (2011) The NLRC4 inflammasome receptors for bacterial flagellin and type III secretion apparatus. Nature. 477, 596–600. doi: 10.1038/nature10510.

Zygiel, E.M., Obisesan, A.O., Nelson, C.E., Oglesby, A.G., Nolan, E.M. (2021). Heme protects Pseudomonas aeruginosa and Staphylococcus aureus from calprotectin-induced iron starvation. J Biol Chem. 296:100160. doi: 10.1074/jbc.RA120.015975.

