## Supplementary material for "The regulatory protein ChuP connects heme and siderophore-mediated iron acquisition systems required for *Chromobacterium violaceum* virulence"

[1]Departamento de Biologia Celular e Molecular e Bioagentes Patogênicos, Faculdade de Medicina de Ribeirão Preto, Universidade de São Paulo, Ribeirão Preto, SP, Brazil

***Correspondence:**

José F. da Silva Neto

### Table S1. Primers used in this work

| Primer name | Sequence (5'→3')[a] | Description |
|---|---|---|
| **Construction of mutant strains** | | |
| chuP_del1[b] | taccggaagcttcccagctgtagtcgatgatg | *Hind*III/*Bam*HI upstream flanking fragment with 656 bp |
| chuP_del2[b] | taccggggatccgggcggtgagtgatatgtgc | |
| chuP_del3 | taccggggatccatcaagtaacacccgcaagcc | *Bam*HI/*Eco*RI downstream flanking fragment with 599 bp |
| chuP_del4 | taccgggaattcttgacgctgccgcctatccc | |
| chuR_del1 | taccggaagcttcccttgcgtgtttcctgcgc | *Hind*III/*Bam*HI upstream flanking fragment with 628 bp |
| chuR_del2 | taccggggatccgacctggatggtctccagcg | |
| chuR_del3[3] | taccggggatcccagaccgccggtttcgagac | *Bam*HI/*Eco*RI downstream flanking fragment with 672 bp |
| chuR_del4 | taccgggaattcttgtgcacggcttctccgtg | |
| chuS_del1 | taccggaagcttagtaccagaacatcggccgg | *Hind*III/*Bam*HI upstream flanking fragment with 642 bp |
| chuS_del2[4] | taccggggatcccagctcgcattcgctgacgc | |
| chuS_del3 | taccggggatccgagcgggcggaatgggtgaa | *Bam*HI/*Eco*RI downstream flanking fragment with 632 bp |
| chuS_del4[6] | taccgggaattcccagcggcttgaaaccgttg | |
| chuTUV_del1 | taccggaagcttagcctgaacgacgtgcacgc | *Hind*III/*Bam*HI upstream flanking fragment with 634 bp |
| chuTUV_del2 | taccggggatcccagctcggcttccttgctgc | |
| chuTUV_del3[b] | taccggggatccaagaccgtcgcccaggtgct | *Bam*HI/*Eco*RI downstream flanking fragment with 678 bp |
| chuTUV_del4[b] | taccgggaattcgcgaagaatacccgctggtg | |
| M13_FW | gtaaaacgacggccagt | Sequencing of cloned fragments |
| M13_RV | agcggataacaatttcac | |
| **Construction of complemented strains** | | |
| chuP_CompFW[c] | taccggggtacctccccttgcgtgtttcctgc | *Kpn*I/*Eco*RI 631 bp fragment with *chuP* and its promoter |
| chuP_CompRV | taccgggaattctgacctggatggtctccagc | region |
| chuR_CompFW | taccggggtaccagtaacacccgcaagccagc | *Kpn*I/*Eco*RI 2423 bp fragment with *chuR* |
| chuR_CompRV | taccgggaattcaccagctcgcattcgctgac | |
| chuS_CompFW | taccggggtaccccaattctgatccacccggc | *Kpn*I/*Eco*RI 1195 bp fragment with *chuS* |
| chuS_CompRV | taccgggaattccgactacgatctcggcgatg | |
| chuTUV_CompFW[5] | taccggggtaccagcgggcggaatgggtgaag | *Kpn*I/*Eco*RI 2633 bp fragment with *chuTUV* |
| chuTUV_CompRV[c] | taccgggaattcctacgcgctgaagtaaggcg | |
| **Heterologous expression** | | |

| | | |
|---|---|---|
| chuP_ExpFW | taccggcatatgagcacatatcactcaccgc | *Nde*I/*Bam*HI 174 bp fragment with *chuP* open reading frame |
| chuP_ExpRV | taccggggatccttacttgatcagtatcagtttgcc | |
| T7_promoter | taatacgactcactataggg | Sequencing of cloned gene |
| T7_terminator | gctagttattgctcagcgg | |
| **β-galactosidase assay and EMSA** | | |
| chuP_promot_FW[d] | taccgggaattcgcttgccgaaggtcagcg | *Eco*RI/*Hind*III 564 bp fragment with *chuP* promoter region for pRK*lacZ*290 cloning |
| chuP_promot_Rv | taccggaagcttgctgcaggcggtagatctc | |
| chuR_promot_FW[1] | taccgggaattctgagcacatatcactcaccg | *Eco*RI/*Hind*III 439 bp fragment upstream of *chuR* for pRKlacZ290 cloning |
| chuR_promot_RV[2,d] | taccggaagcttttcgccgcgtcggcaatatc | |
| CV_2599_promotFW | cctagcgaattcgcggccaaagagtcaggaa | *Eco*RI/*Bam*HI 288 bp fragment with CV_2599 promoter region |
| CV_2599_del2 | ggcctaggatcctacccgttgtacggcagcg | |
| l*acZ*290up | tgacggctatcaccatca | Confirmation of cloned sequences into pRK*lacZ*290 |
| **RT-qPCR** | | |
| CV_3896NB_FW | gaccttccccagcaagaccttc | 124 bp fragment of *chuR* coding region |
| CV_3896NB_RV | cttgtaatggtcccagcgcagc | |
| vbaF_RT-qPCR_FW | cgctgcagtacggactgga | 110 bp fragment of *vbaF* coding region |
| vbaF_RT-qPCR_RV | tatggtgtcggcgccatac | |
| vbuA_RT-qPCR_FW | cctgaacatgcgtttggacg | 122 bp fragment of *vbuA* coding region |
| vbuA_RT-qPCR_FW | gcaggttttggtcacgatcg | |
| CV_4206NB_FW | gaaaaaccgctcttcatgctg | 122 bp fragment of *rpoH* coding region |
| CV_4206NB_FW | gatcgtggttgccgctgaaa | |

[a] Digestion sites recognized by the restriction enzymes are underlined.

[b]These primers were also used to delete the entire *chu* operon.

[c]These primers were also used to complement the entire *chu* operon.

[d] These primers were also used to generate the p*chuPR* EMSA probe.

[1-6]Primers employed in RT-PCR.

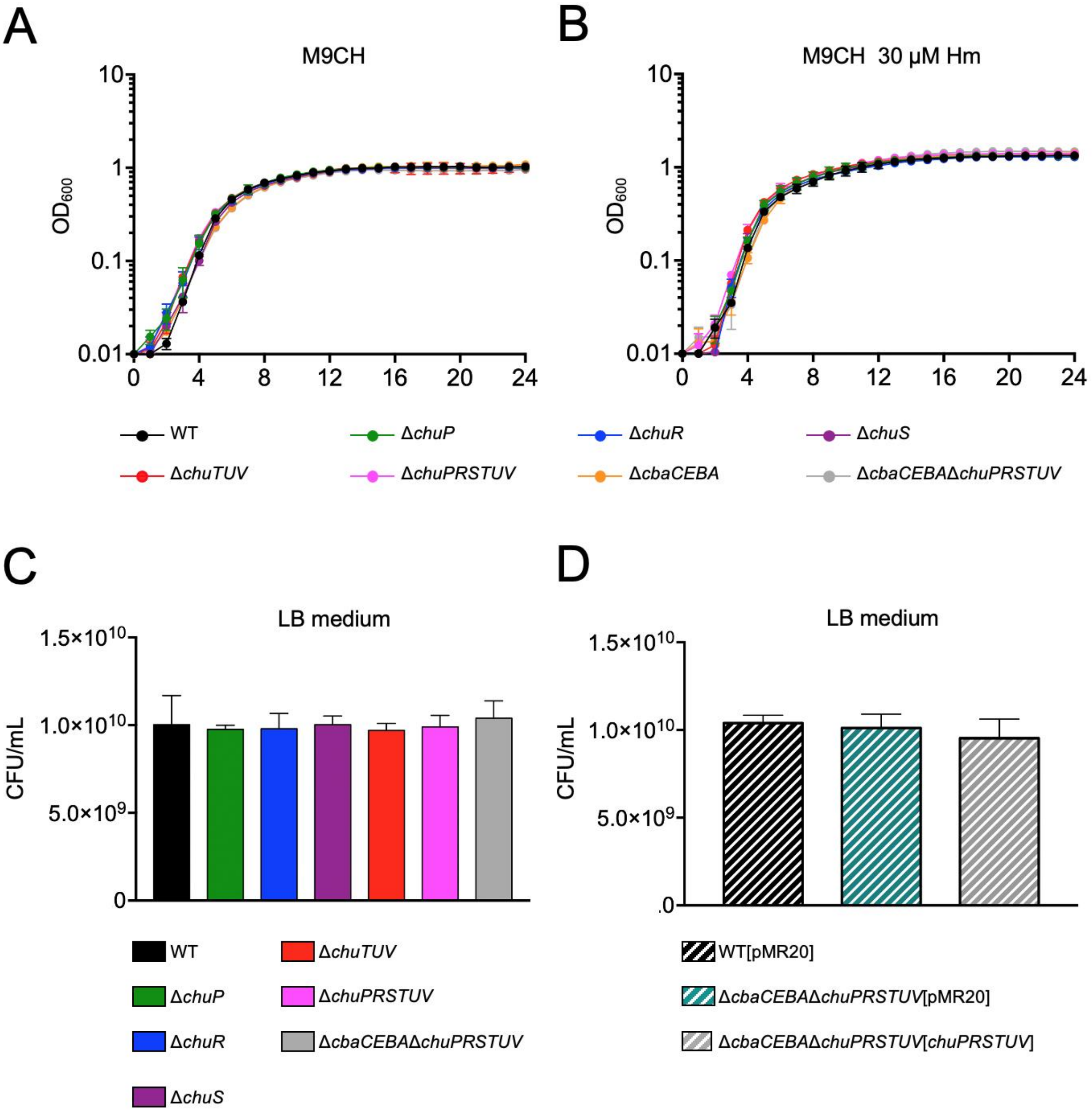


**Supplementary Figure 1.** Mutants of the *chu* operon have regular fitness under standard growth conditions. **(A and B)** Growth curves. The WT and the indicated mutant strains were grown in M9CH without **(A)** or with Hm supplementation **(B)** from an $OD_{600}$ of 0.01. The $OD_{600}$ was measured every 15 minutes for 24 h. Data points are shown in 1-h intervals as the mean and standard deviation of three biological replicates. **(C and D).** Cell viability in LB medium by CFU counting. The indicated mutant **(C)** and complemented **(D)** strains were grown for 20 h in LB, serial diluted, and plated for CFU quantification. Data are from three biological replicates. Mutant and complemented strains were compared to WT and WT[pMR20], respectively. When not indicated, n.s. (not significant). One-way ANOVA followed by Tukey's multiple-comparison test.

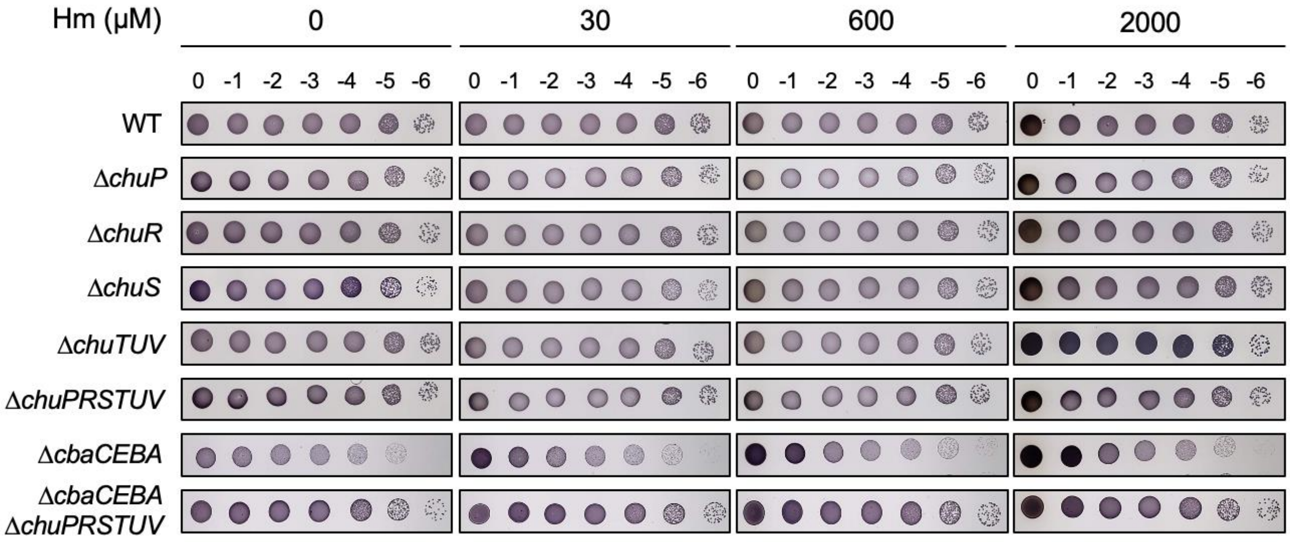


**Supplementary Figure 2.** Mutants of the *chu* operon have regular fitness under heme excess. *C. violaceum* WT and the indicated mutant strains were grown from an $OD_{600}$ of 0.01 in either M9CH or M9CH supplemented with the indicated Hm concentrations. After 24 h cultivation, the cultures were serial diluted and platted on M9CH to evaluate cell viability. Data representative of three biological replicates.

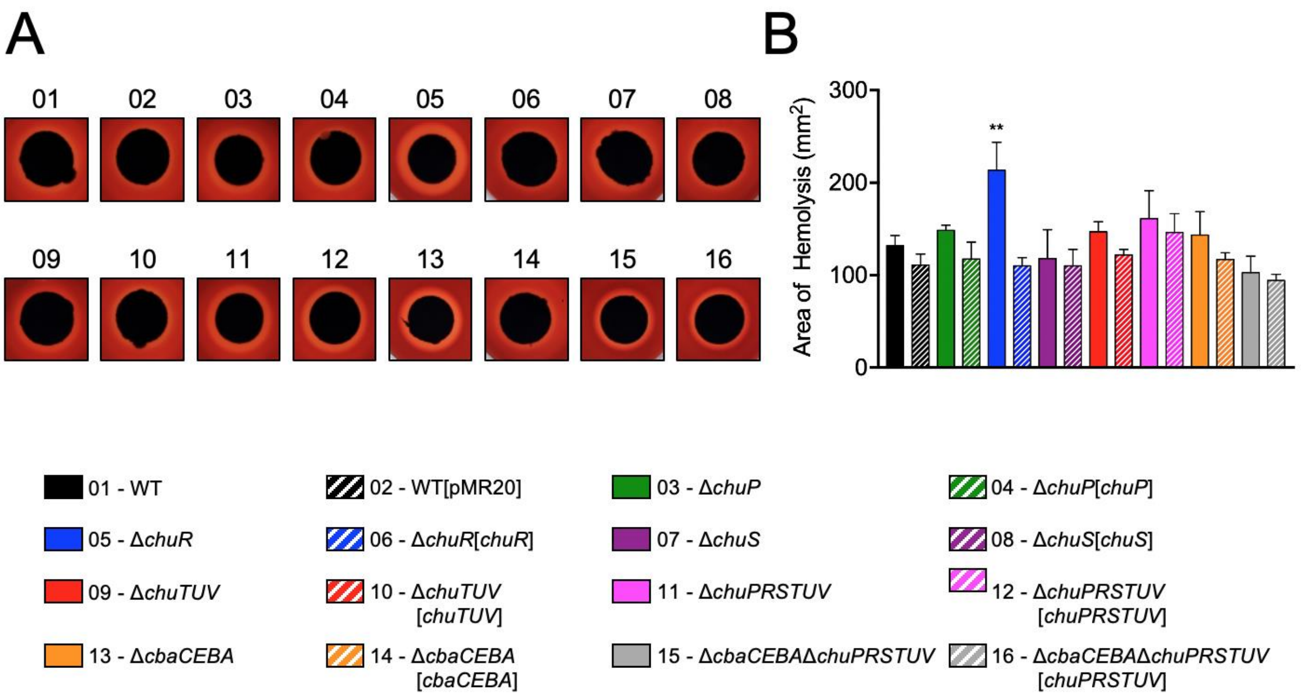


**Supplementary Figure 3.** Deletion of *chuR* increases hemolytic activity. **(A)** Hemolysis in blood agar. The indicated strains were grown in M9CH and spotted on 5% (v/v) sheep-blood Mueller-Hinton plates. The lighter halos around bacteria growth indicate hemolysis. Representative images of one assay are shown. **(B)** Quantification of hemolytic activity. The area of the lighter halos was measured using Image J software, subtracting the area of bacterial growth to eliminate differences in sizes. Data are from three biological replicates. Mutant and complemented strains were compared to WT and WT[pMR20], respectively. **, $p < 0.01$; when not shown, n.s. (not significant). One-way ANOVA followed by Tukey's multiple-comparison test.
